# Manganese Homeostasis Drives *Stenotrophomonas maltophilia* Oxidative Stress Defense and Replication in *Acanthamoeba castellanii* phagosomes

**DOI:** 10.1101/2025.10.14.682371

**Authors:** Fulvia-Stefany Argueta-Zepeda, Javier Rivera, Julio César Valerdi-Negreros, Christopher Rensing, Pablo Vinuesa

## Abstract

Manganese homeostasis is essential for the environmental adaptability and pathogenic potential of *Stenotrophomonas maltophilia*, a bacterium that thrives across diverse and fluctuating niches. Here, we characterize the manganese homeostasis network of *S. maltophilia* strain Sm18, revealing a coordinated system that integrates conserved transporters with previously unrecognized components. Central to this system is an MntR-controlled miniregulon that includes the canonical Mn²⁺ importer MntH and exporter MntP, together with a TonB-dependent receptor (TBDR) and a periplasmic thioredoxin-fold protein (pTFP), both defining novel protein families with restricted phylogenetic distribution. Transcriptomic analyses under varying Mn²⁺ and Fe²⁺ conditions uncovered a tight interplay between these metals, highlighting the ferrophilic nature of *S. maltophilia* and the differential regulation of miniregulon components. Notably, the TBDR-pTFP module is strongly induced under combined Mn^2+^ and Fe^2+^ limitation, suggesting a specialized role in metal acquisition under nutrient-restricted conditions. Functional analyses demonstrated that MntP is required to prevent Mn toxicity even at sub-inhibitory concentrations, whereas MntH supports growth under oxidative stress and promotes intracellular replication within *Acanthamoeba castellanii* phagosomes. Together, these findings define a previously unrecognized Mn-responsive module that expands the MntR regulatory network and provides new insight into the mechanisms that enable *S. maltophilia* to adapt to metal-limited and host-associated environments.

## 1. Introduction

*S. maltophilia* is a gram-negative, non-fermentative bacterium recognized for its ecological versatility and opportunistic pathogenicity (Brooke, 2021). It thrives in diverse environments, including soil, water, plant rhizospheres, free-living amoebae, and diverse animal organs and cells, demonstrating remarkable adaptability to contrasting ecological niches (Ochoa-Sánchez and Vinuesa, 2017; Rivera et al., 2024; Ryan et al., 2009). These bacteria can survive harsh conditions, such as oligotrophic or metal-rich environments, and are highly resistant to disinfectants and multiple drugs due to robust biofilm formation and multiple efflux systems. Recent research has highlighted these adaptive mechanisms, showing how they enhance the resilience and persistence of *S. maltophilia* in nosocomial environments (Brooke, 2021, 2012), making it one of the ten most prevalent microorganisms in hospital care units (Rello et al., 2019). This pathogen primarily affects immunocompromised individuals, leading to respiratory tract, bloodstream, and urinary tract infections (Brooke, 2021), and poses a significant challenge in clinical settings because of its intrinsic resistance to multiple antibiotics, including carbapenems (Chang et al., 2015; Ochoa-Sánchez and Vinuesa, 2017). However, little is known about the genes and mechanisms involved in the adaptation of *Stenotrophomonas* to host-associated and intracellular lifestyles in diverse host tissues and phagocytic cells, including free-living amoebae (Denet et al., 2018; Rivera et al., 2024) and mammalian macrophages (Takahashi et al., 2020). In recent years, mounting evidence has revealed that professional phagocytes subject ingested bacteria to limiting or toxic concentrations of transition metals, inhibiting their proliferation in phagosomes, a process known as nutritional immunity (Murdoch and Skaar, 2022).

Homeostasis of essential transition metals, such as manganese, iron, copper, and zinc, is crucial for various physiological processes (Palmer and Skaar, 2016; Waldron and Robinson, 2009). Metallostasis is maintained by a network of allosteric metalloregulators, metal transporters, metallochaperones, and efflux pumps (Porcheron et al., 2013). The concerted action of these components regulates the intracellular metal concentrations, ensuring that they are available for essential functions without reaching toxic levels (Chandrangsu et al., 2017). The ability of bacteria to rapidly adapt to different environments is tightly coupled to their metal homeostasis capabilities, particularly during host infection, as metal concentrations vary widely across organs, tissues, and cellular compartments (Kehl-Fie et al., 2013; Pompilio et al., 2014).

Studies on *S. maltophilia* metallostasis have focused on iron homeostasis, highlighting its importance in bacterial virulence (Mikhailovich et al., 2024). In contrast, our knowledge of the molecular mechanisms underlying manganese homeostasis in *Stenotrophomonas* is limited to a single transcriptomic study performed on *Stenotrophomonas* sp. MNB17, an organism isolated from a deep-sea mud sample in the western Pacific that removes Mn^2+^ by efficiently forming Mn-precipitates (MnCO_3_ and Mn oxides) at high Mn concentrations (Song et al., 2022). However, manganese homeostasis has been well studied in some Gram-positive and Gram-negative bacteria (Bosma et al., 2021; Waters, 2020). A crucial component is the manganese transporter protein MntH, which facilitates manganese uptake into the cytoplasm under limiting conditions (Kehres et al., 2000). Conversely, the conserved MntP exporter removes excess manganese, thereby maintaining intracellular levels within the optimal range (Waters et al., 2011). This process is regulated by various regulatory proteins and signaling pathways that sense intracellular manganese levels and adjust the expression or activity of manganese transporters (Bosma et al., 2021; Martin and Waters, 2022; Sengupta et al., 2018; Wright et al., 2024), with MntR being the master regulator (Que and Helmann, 2000; Waters et al., 2011). Furthermore, under manganese limitation and oxidative stress, a Mn^2+^-scavenging pathway has been identified in *Burkholderia thailandensis*, comprising a newly discovered TonB-dependent outer membrane (OM) manganese transporter, MnoT, and a Mn^2+^-binding protein, TseM, secreted by a type VI secretion pathway (T6SS) which is apparently unique to *Burkholderia* species (Si et al., 2017). The requirement for Mn^2+^ and Fe^2+^ varies greatly among bacterial species (Bosma et al., 2021). *S. maltophilia* features an iron-centric metabolism that requires iron for optimal growth and virulence (Kalidasan et al., 2018; Mikhailovich et al., 2024). However, Mn^2+^ becomes essential under oxidative stress or low iron availability (Anjem et al., 2009), conditions that mirror the environments that *S. maltophilia* likely encounters as an intracellular opportunistic pathogen in phagosomes subjected to nutritional immunity (Čapek and Večerek, 2023; Murdoch and Skaar, 2022). This bacterium has been reported to survive in human epidermoid epithelial cells (Zhang et al., 2024), human lung epithelial cells (DuMont et al., 2015; Nas et al., 2019), free-living amoebae (Denet et al., 2018; Rivera et al., 2024), and mammalian macrophages (Nas et al., 2019; Takahashi et al., 2020). The divalent Mn^2+^/Fe^2+^ transporter Nramp1 protein, located in the phagosomal membrane and ubiquitous in eukaryotes, plays a crucial role in nutritional immunity by depleting phagocytosed bacteria of these micronutrients. Interestingly, its bacterial homolog, MntH, is also essential for countering this stress by importing Mn^2+^ into bacterial cells (Cellier, 2001; Juttukonda and Skaar, 2015).

Here, we investigated the manganese homeostasis network of*S. Maltophilia* Sm18 to address the limited understanding of this process in the genus. Through an integrated transcriptomic, phylogenomic and functional approach, we identified the core components of an MntR-controlled miniregulon, including the canonical transporters *mntH* and *mntP*, as well as a previously uncharacterized bicistronic locus encoding a TonB-dependent receptor and a thioredoxin-fold protein. We further assessed the contribution of these components to bacterial fitness under metal stress conditions and within a biologically relevant intracellular context. In particular, we demonstrate that *mntH* is expressed during interaction with *A. castellani*i and is required for optimal intracellular replication, linking manganese acquisition to adaptation to host-associated environments.

## 2. Material and methods

### 2.1 Bacterial strains and culture conditions

*S. maltophilia* ESTM1D_MKCAZ16_6a (Sm18) is an environmental isolate from river sediments in Morelos, Mexico (Ochoa-Sánchez and Vinuesa, 2017) with a fully sequenced genome (Vinuesa et al., 2026c). Sm18 was routinely cultivated aerobically in Lysogeny Broth (LB) at 30°C or in modified MOPS minimal medium (MM) supplemented with.4 % (w/v) glucose (LaBauve and Wargo, 2012). The MM was optimized for *S. maltophilia* growth by adding thiamine 0.5%, biotin 0.1% and 0.2% (w/v) casamino acids (CAA). Manganese was not included in the micronutrient stock of MM; however, trace amounts of metals, including iron and potentially manganese, may be present due to the addition of casamino acids, as indicated by the manufacturer. The addition of casamino acids was required to support robust growth of Sm18 strains carrying transcriptional fusions, which did not grow reproducibly in MOPS minimal medium supplemented only with methionine. For the growth kinetics, LB was supplemented with 10mM of MOPS buffer (pH 6.8) and antibiotics, as required.

### 2.2 Generation of deletion and vector integration mutants

Markless deletion mutants were generated using a two-step allelic exchange strategy (Huang and Wilks, 2017). Briefly, ∼ 500-600 bp DNA fragments flanking the target genes were PCR-amplified from *S. maltophilia* Sm18 genomic DNA and assembled into the suicide vector pEX18Tc (Hoang et al., 1998) using NEBuilder HiFi DNA Assembly (NEB), following the manufacturer’s instructions. The resulting constructs were transferred into Sm18 by conjugation from *E. coli* S17-1, as previously described (Simon et al., 1983). Transconjugants (merodiploids) were selected on LB agar supplemented with tetracycline (20 µg/mL) and gentamicin (30 µg/mL). Double recombinants were obtained by counter-selection on LB agar without NaCl and supplemented with 10% sucrose. Markerless deletions were confirmed by antibiotic sensitivity, colony PCR, and amplicon sequencing. This approach was used to generate deletion mutants of *orf02357*, *orf02358*, *mntP* and *mntR*. For *mntH*, repeated attempts to obtain a deletion mutant were unsuccessful. Therefore, a vector integration mutant (VIM) strategy was used to disrupt gene function. Briefly, an internal fragment of the target gene was PCR-amplified and cloned into pEX18TcVIM-GFP, followed by conjugation into Sm18 as described above. This approach was also applied to selected genes (*mntP* and *orf02357*) to validate that VIM and deletion mutants produced comparable phenotypes (Supplementary Fig. S1). All primers and constructs used in this study are listed in Supplementary Tables A4-A6.

### 2.3 Construction of Sm18 expression plasmids

Expression plasmids were constructed based on the standardized SEVA modular architecture (Martínez-García et al., 2023). Derivatives of pSEVA327 (oriVRK2, CmR) and pSEVA337R (oriVpBBR1, CmR) were used to generate transcriptional reporter fusions. Intergenic regions upstream of *orf02357*, *mntH,* and *mntP* were PCR-amplified from Sm18 genomic DNA using Phusion polymerase (NEB). The 5’-end UTR found upstream of *orf2357* also controls the transcription of *orf2358*, which is co-expressed with the preceding gene and separated by a short intergenic distance. Amplicons were directionally cloned upstream of promoterless GFP or mCherry reporter genes using standard restriction-ligation procedures. The resulting constructs (327-02357_02358, 327-*mntH,* 327-*mntP,* 337R-02357_02358, 337R-*mntH,* and 337R-*mntP*) are listed in Supplementary Tables A4-A6.

### 2.4 Fluorescence kinetics and microbial growth

Fluorescence and growth kinetics of strains were assessed in at least five and eight independent biological replicates, respectively, using 96-well microplates. Cultures were adjusted to an initial OD_600_ of 0.05 in LB and 0.01 in minimal MOPS (MM) medium, in a final volume of 200 µL supplemented with varying Mn and Fe concentrations. Plates were incubated at 30 °C with orbital shaking (205 rpm), and OD_600_ measurements were recorded every 30-45 min using a BioTek Epoch 2 microplate spectrophotometer. Fluorescence (GFP; excitation/emission: 479/520 nm) and OD_600_ were measured in parallel under identical conditions using a BioTek Synergy H1 microplate reader. Uninoculated media were used as blanks, and strains carrying empty plasmids served as controls. Relative fluorescence units (RFU) were normalized to OD_600_ previous statistical analysis. Growth and fluorescence kinetics were analyzed using the area under the curve (AUC), calculated with the Growthcurver R package (v0.3.1). Statistical comparisons between conditions were performed using the non-parametric Kruskal-Wallis test followed by Dunn’s post hoc test with Bonferroni or Benjamini-Hochberg correction for multiple testing. Bootstrapping was used to estimate 95% confidence intervals. All analyses were conducted in R (v4.3.2) using standard packages (ggplot2, ggpubr, rstatix). Error bars represent the standard error of the mean from biological and technical replicates.

### 2.5 Cell sample preparation for RNA-sequencing

Three independent biological replicates were generated from glycerol stocks by streaking on LB agar and incubating at 30 °C. A single colony from each replicate was inoculated into 3 mL of MM and grown at 30 °C with shaking (205 rpm) for 24 h. Precultures (1:50 dilution) were then established in 25 mL MM and incubated under the same conditions for 14 h. Cells were harvested by centrifugation, washed three times with MM without added iron (MM-Fe0), and used to inoculate fresh MM at an initial OD_600_ of ∼0.005. Cultures were grown in media supplemented with defined concentrations of Mn^2+^ and Fe^2+^, denoted as Fe0Mn0, Fe0Mn8, Fe10Mn0, and Fe10Mn8, where the numbers indicate the micromolar concentration (µM) of each metal added (e.g., MM-Fe0Mn8 indicates 0 µM Fe^2+^ and 8 µM Mn^2+^). Fe0 and Mn0 refer to conditions without added metals; however, trace amounts may still be present due to the composition of the basal medium (see Section 2.1). All experimental conditions were prepared using the same basal medium, such that relative differences in gene expression and phenotypes reflect the defined supplementation of Fe^2+^ and Mn^2+^. Cultures were grown to early exponential phase (OD₆₀₀ 0.1–0.2), then split into two technical replicates and harvested by centrifugation. Cell pellets were immediately stabilized in RNA later for RNA sequencing (GENEWIZ, Azenta Life Sciences; Illumina HiSeq 2500, 150 bp paired-end).

### 2.6 RNA-seq analysis data and gene expression

Raw reads were quality-checked using FastQC (v0.12.1) (Andrews, S., 2010) and trimmed with Trimmomatic (v0.39) (Bolger et al., 2014). Clean reads (∼135 bp; >23 million paired-end reads per sample) were aligned to the *S. maltophilia* Sm18 reference genome (Vinuesa et al., 2026c) using Bowtie2 (v2.4.4) (Langmead and Salzberg, 2012), achieving alignment rates above 96%. Gene-level counts were obtained using featureCounts from the Rsubread package (v2.14.2) (Liao et al., 2019), with >92% of reads successfully assigned to annotated features. Differential gene expression analysis was performed using DESeq2 (v1.42.0) (Love et al., 2014), applying median-of-ratios normalization and a negative binomial model.

Adjusted p-values were calculated using the Benjamini-Hochberg method, and genes with Padj < 0.01 and |log₂ fold change| ≥ 1 were considered differentially expressed. Functional annotation of differentially expressed genes was performed using UniProt (The UniProt Consortium, 2017), InterProScan (Blum et al., 2025), and MOTIF search tools.

### 2.7 Identification of transcription factor binding sites and protein homologs

Putative MntR-binding motifs were identified using MEME (v 5.5.5) (Bailey et al., 2015) on the 5’-UTR regions of *Stenotrophomonas mntH* (V8P27_002360) orthologs identified with GET_HOMOLOGUES (Contreras-Moreira and Vinuesa, 2013). Motifs ranging from 16 to 25 nt were searched, and the resulting position-specific probability matrices (PSPMs) were compared against experimentally validated transcription factor-binding sites, using TOMTOM with the CollecTF (Erill, 2015) and PRODORIC (Dudek and Jahn, 2022) database. Motif occurrences were further identified using MAST across representative genomes of *Stenotrophomonas* and related genera in the *Lysobacteriaceae* (Chauviat et al., 2025; Vinuesa et al., 2018). Homologs of MntR miniregulon components were identified by hmmsearch (v3.4) (44) using the gathering threshold (--cut_ga) option for calibrated profile hidden Markov models (HMMs) retrieved from Pfam (PF01566 for MntH, PF02659 for MntP), or built locally with hmmbuild (v3.4) from clustal omega (Sievers and Higgins, 2018) alignments (with –iter 2 and –use-kimura options) of our curated datasets of MntR, V8P27_002357, and V8P27_002358 homologs. The latter three HMMs and the PSPM for the *Lisobacteraceae* MntR-box are available for download from https://github.com/vinuesa/supplementary_materials/.

### 2.8 Phylogenomic and motive distribution analysis

A dataset of 103 RefSeq genomes representing the taxonomic diversity of *Stenotrophomonas* and related Lysobacteraceae was compiled. Homologs of MntR, MntH, MntP, ORF2357, and ORF2358 were identified using hmmsearch (HMMER3) with curated profile HMMs. Motif searches for the MntR-binding site were performed using MAST (MEME suite) with the consensus motif defined in this study with a *p* ≤ 5×10^−8^ cutoff. Phylogenetic relationships were inferred using a maximum likelihood approach based on a concatenated alignment of 155 core proteins identified with GET_PHYLOMARKERS (Vinuesa et al., 2018) from a set of 601 core gene families generated with GET_HOMOLOGUES. Presence-absence patterns of genes and motif counts were mapped onto the resulting phylogeny.

### 2.9 Structural and phylogenomic characterization of miniregulon-associated proteins

Protein domain architecture and signal peptides were predicted using standard bioinformatic tools. Structural models were generated using AlphaFold3 (Abramson et al., 2024) and visualized to uncover potential ligand-binding and other functional features. Homologs of ORF2357 and ORF2357 were identified using BLASTP and HMMER searches against public databases, including RefSeq and Reference Proteomes. Retrieved sequences were aligned using Clustal Omega, and phylogenetic relationships were inferred using maximum likelihood methods as described above. Synteny analysis was performed by examining the genomic context of representative homologs across selected taxa. Conserved sequence motifs were identified from multiple sequence alignments and visualized accordingly.

### 2.10 Live-cell imaging (LCI) of of Sm18 expressing transcriptional fusions in co-culture with A. castellanii

Live-cell imaging assays were performed as previously described (Rivera et al., 2024). Briefly, 5 x 10^5^ trophozoites of *A. castellanii* strain Neff were seeded in 20-mm glass-bottom dishes and incubated for 1 h at 30 °C in MMsalts-MOPS-Glc medium. *S. maltophilia* Sm18/337R-*02357_02358*, Sm18/337R-*mntH* and Sm18/337R-*mntP* were grown in LB, washed, and adjusted to an OD_600_ of ∼1.0 (∼4.8 × 10^8^ CFU/mL). Bacterial suspensions were added to the amoeba cultures at a multiplicity of infection (MOI) of 25, followed by centrifugation at 500 × g for 5 min to synchronize phagocytosis. Live-cell imaging was performed using an AxioVert A1 inverted microscope (Zeiss) at 3 and 18 h post co-culture (ppc).

### 2.11 Quantitative evaluation of the intracellular replication capacity of Sm18 mutants

Intracellular replication assays were performed using a microtiter plate format as previously described (Rivera et al., 2024). Briefly, 1 x 10^5^ trophozoites of *A. castellanii* (strain Neff) were seeded per well in MMsalts-MOPS-Glc medium and incubated for 30 min at 25° C. The medium was then replaced with Sm18VIM*mntH* and Sm18Δ*mntR* bacterial suspensions tagged with the minitransposon pUC18T_mTn7TC1_Pc_mScarlet-I, which constitutively expresses the mScarlet-I fluorescent protein from strong Pc promoter, at a MOI of 10.

Phagocytosis was synchronized by centrifugation and plates were incubated statically at 25 °C in a Synergy 2 plate reader. Red fluorescence (excitation/emission: 568/594 nm) was measured every 30 min for 48 h. Control wells included medium only, trophozoites without bacteria, and bacteria without trophozoites. All experiments were performed with five independent biological replicates per strain, each including five technical replicates.

## 3 RESULTS

### 3.1 Manganese tolerance of S. maltophilia Sm18 in complex and defined media

To assess the strain’s manganese tolerance, we ran 48-hour-long growth kinetics using LB and MM media with increasing MnCl_2_ concentrations. Strain Sm18 grew in LB and MM up to 16 mM and 8 mM of MnCl_2_, respectively (Fig. 1A). In both media, the maximal growth rate and total yield of the culture remained unaffected, except at the highest Mn concentration tested. However, statistical analysis of the area under the curve (AUC) for population density (OD_600_) data revealed a significant decrease in growth in both media starting at 8 mM MnCl_2_, due to the extended lag phase observed with increasing Mn concentrations (Fig. 1B).

**Fig. 1.**
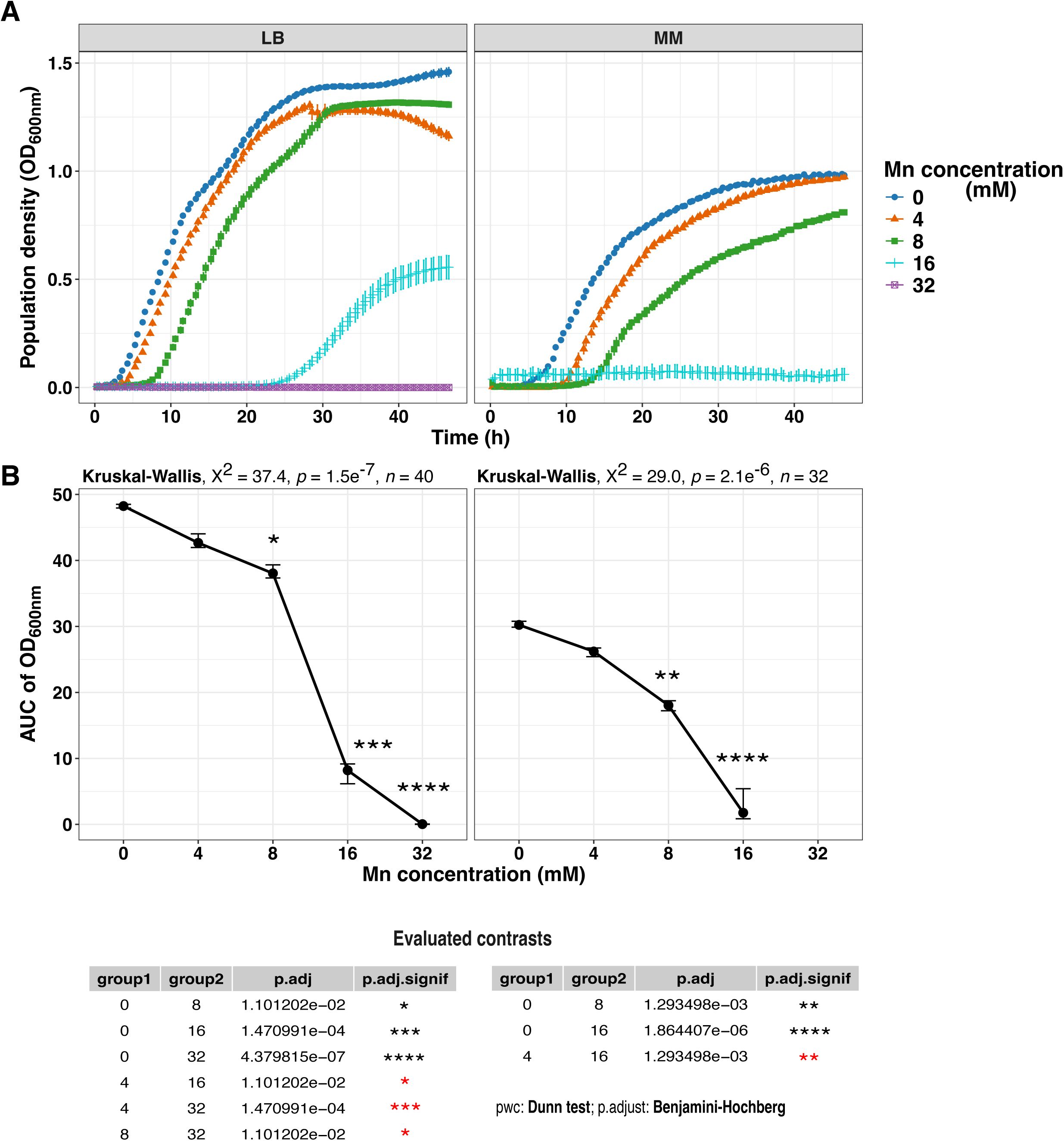
Growth kinetics of *S. maltophilia* Sm18 under increasing Mn concentrations. (A) Growth in Lysogeny Broth (LB) and minimal MOPS (MM) medium with increasing millimolar (mM) concentrations of manganese (Mn^2+^). The means of eight biological replicates per concentration are represented by distinct shapes and colors, with error bars indicating the standard error. (B) Statistical analysis of Mn tolerance in Sm18. Differences among groups were assessed using the non-parametric Kruskal-Wallis test, followed by Dunn’s *post hoc* test with Benjamini-Hochberg (BH) correction. Significance levels are shown as: ns, *p* > 0.05; *, *p* < 0.05; **, *p* < 0.01; ***, *p* < 0.001; ***, *p* < 0.0001. Black asterisks indicate comparisons with the manganese-free control, while red asterisks indicate comparisons between other groups.

Consistent with these observations, complete growth inhibition was observed at higher Mn^2+^ concentrations, occurring at 32 mM in LB and at 16 mM in MM medium. Although minimum inhibitory concentrations (MICs) were not determined using standardized assays, these results define the concentration range at which Mn²⁺ becomes inhibitory under the tested conditions. The lower inhibitory thresholds were observed consistently across replicates.

### 3.2 Genome-wide identification of manganese-responsive genes in S. maltophilia Sm18

To identify Sm18 genes involved in Mn^2+^ hemostasis, we designed experimental conditions that consider the well-established interplay between iron and manganese in bacterial metal homeostasis (Čapek and Večerek, 2023). These metals are known to be functionally interconnected, often sharing regulatory networks and compensatory roles under conditions of metal limitation (Helmann, 2014; Puri et al., 2010; Steingard et al., 2023; Taudte et al., 2016). Therefore, varying iron availability alongside manganese allows a more accurate assessment of manganese-dependent regulatory responses. Cultures were established in defined MM medium supplemented with casamino acids under four metal conditions: Fe0Mn0, Fe0Mn8, Fe10Mn0, and Fe10Mn8 (see methods). Principal component analysis (PCA) shown in Fig. 2A demonstrated that replicates within each group were clustered and separated according to metal concentration. PC1 revealed that 64% of the variance was related to varying iron conditions, while PC2 explained 10% of the variance due to manganese concentrations. We identified a total of 124 differentially expressed genes (DEGs) when comparing Fe^2+^ 10 µM vs. Fe^2+^ 0 µM in MM-Mn0, and 89 DEG in MM-Mn8, based on an adjusted *p*-value (padj) < 0.01 and a log₂ fold change |(log₂FC)| ≥ 1.0. In contrast, only five DEGs were detected when comparing Mn^2+^ 0 µM vs. Mn^2+^ 8 µM in MM-Fe10, and 16 DEG in MM-Fe0. The number of DEGs for each condition is summarized as a Venn diagram (Fig. 2B), revealing that 88 DEGs were regulated solely by iron levels, irrespective of manganese concentrations in the medium. Although this study focuses on manganese-related responses modulated by manganese and iron availability, we identified a broader transcriptional response reflecting the interplay between these two metals. A complementary analysis specifically addressing iron homeostasis will be presented in a separate study. The *mntP* gene was the only one found to be exclusively regulated by manganese levels in the medium, independent of iron availability (Fig. 2B). In contrast, the expression of *mntH* was affected by any change in Mn or Fe concentration (Fig. 2B). Nine genes displayed differential expression only when cultures were subjected to simultaneous Fe^2+^ and Mn^2+^ limitation (Fig. 2B). Two were down-regulated (blue-colored locus tags) under these conditions, whereas the remaining genes were up-regulated, as detailed below. Figure 2C highlights the DEGs identified when comparing Mn^2+^ 0 µM vs. Mn^2+^ 8 µM in both MM-Fe10 and MM-Fe0 media (see Supplementary Data, Tables A1 and A2 for more details). These DEGs are represented as colored dots on the log_2_FC plots, and their corresponding genomic loci are indicated by green vertical bars mapped to the Sm18 chromosome. Their gene ontologies (GOs) are highlighted in different colors with the transmembrane transport category (yellow points) being the most prevalent. Fig. 2C also shows the log_2_FC values in a bar plot for a better quantitative representation, including the protein annotations of each DEG. Loci displaying the most significant log_2_FC changes were selected for further characterization.

**Fig. 2.**
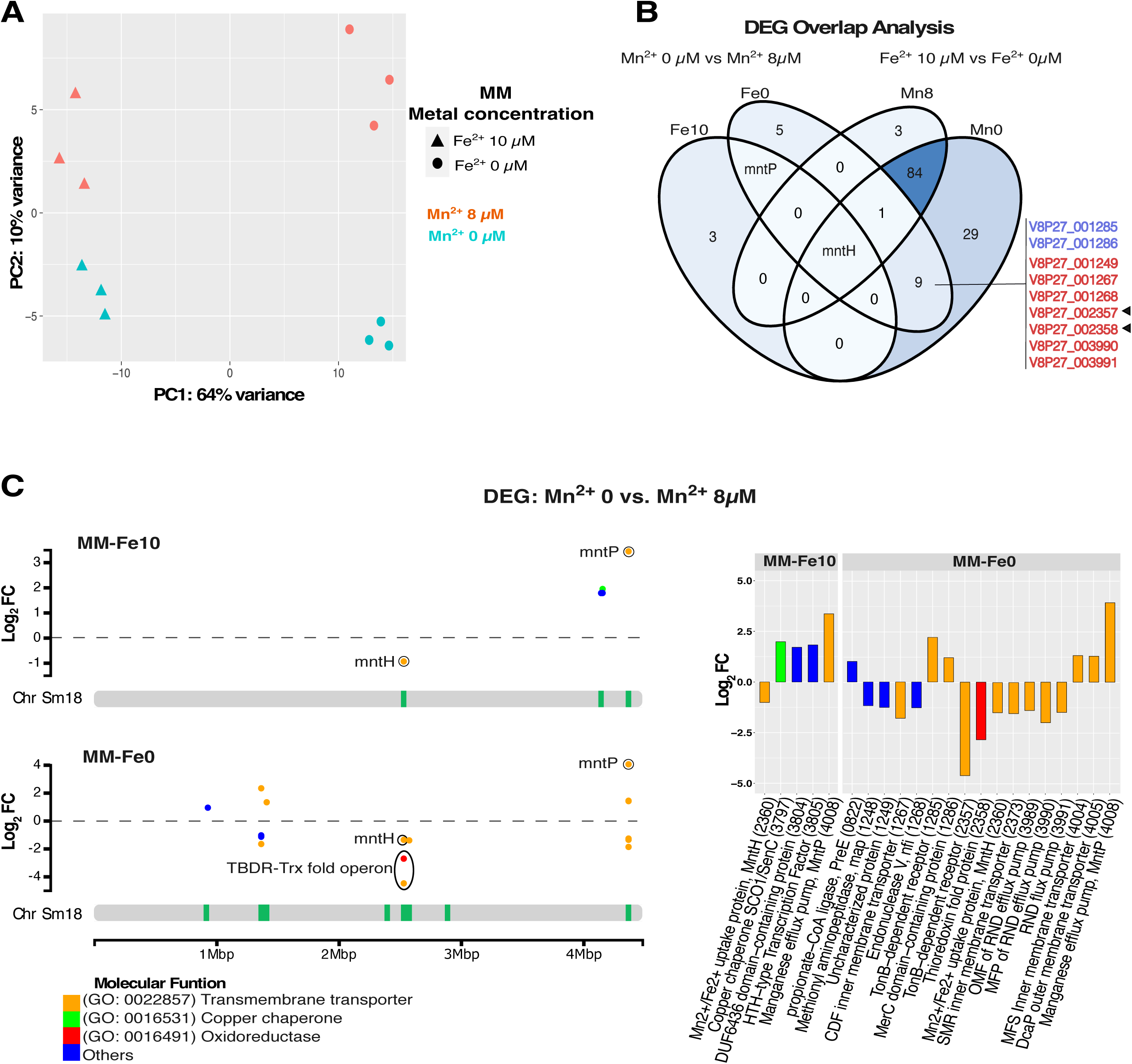
Transcriptomic analysis and identification of differentially expressed genes (DEG). Data were obtained from three biological replicates. (A) Principal component analysis (PCA) based on DESeq2-normalized read counts from the full transcriptome (*n* = 4,059 genes). (B) Venn diagram generated using the VennDiagram package in R, illustrating the overlap of DEG at different metal concentrations. Genes were considered differentially expressed if they met the criteria of adjusted *p*-value (padj) < 0.01 and |log_2_ fold change| ≥ 1. (C) DEG from Mn^2+^ 0 µM vs. Mn^2+^ 8 µM comparisons in MM-Fe10 and MM-Fe0 mapped across the Sm18 chromosome (Chr Sm18). Green loci denote regions with significant differential expression. Colored dots represent genes classified by Gene Ontology (GO) functional categories. The adjacent bar plot displays the protein annotations of each DEG.

### 3.3 Genome-wide scanning for potential MntR binding sites

MntR is a manganese-responsive metalloregulator that controls gene expression by binding to conserved short inverted palindromic sequences (MntR-boxes) located in the 5′-UTR regions of target genes (Patzer and Hantke, 2001). Using a motif derived from *mntH* upstream regions, we identified three MntR-binding sites in the Sm18 genome (Fig. 3A-C). In addition to the canonical target *mntH,* two additional loci were identified as putative members of the MntR miniregulon. One binding site was located upstream of the *orf2357-orf2358* locus, which lies in close proximity to *mntR* and *mntH*, which share a bidirectional promoter (Fig. 3B). Additionally, *orf2357* and *orf2358* are separated by a minimal intergenic region, consistent with their organization as a putative operon (Fig. 3B). A third MntR-binding site was identified upstream of *mntP*, where a putative yybP–ykoY riboswitch was also detected (Fig. 3C). Together, these findings identify a set of three loci associated with predicted MntR-binding sites in the Sm18 genome which also correspond to the genes displaying the highest differential expression in the transcriptomic analysis. We next examined the phylogenetic distribution of these loci, considered here as candidate components of a MntR miniregulon of manganese homeostasis, across the genus *Stenotrophomonas* and related Lysobacteraceae.

**Fig. 3.**
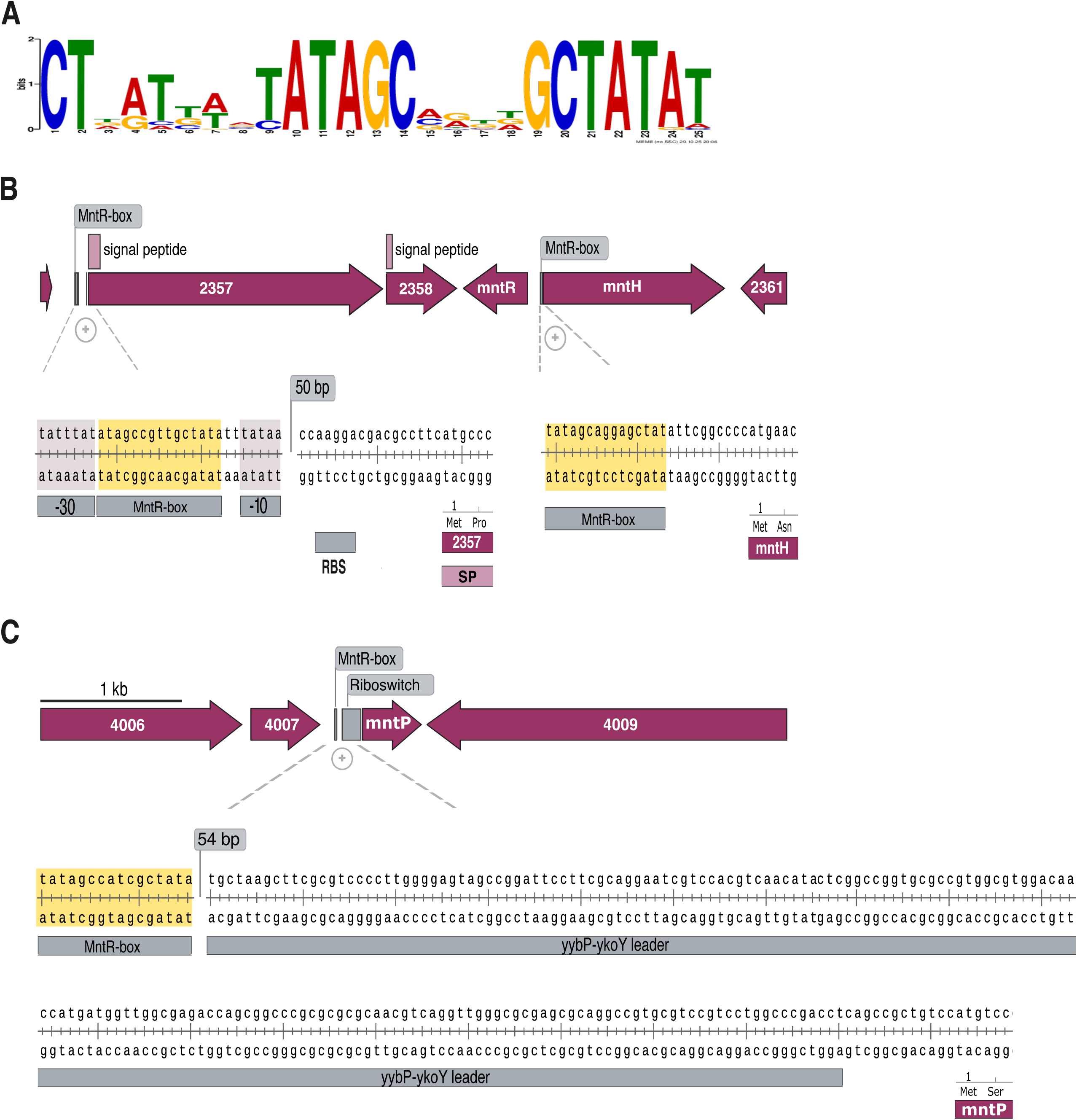
Core components of the MntR miniregulon in *S. maltophilia* Sm18. (A) Binding motif (MntR-box) for the MntR transcription factor identified using MEME-MAST. The MntR-box profile corresponds to a 25-bp inverted palindromic sequence. (B) Putative manganese uptake operon encoding an outer membrane TonB-dependent receptor and a periplasmic thioredoxin-fold protein, identified by locus tags 2357 and 2358, respectively. The MntR-box is located 75 bp upstream of the putative start codon, between the −10 and −35 promoter regions. These genes are adjacent to *mntR* (encoding the transcription factor MntR) and *mntH* (encoding the cytoplasmic Mn importer), which share a 117 bp intergenic region. The mntH MntR-box is located 10 bp upstream of the putative start codon. (C) The gene encoding the inner membrane manganese exporter MntP is predicted to be regulated by both the *yybP-ykoY* riboswitch and MntR. Its MntR-box is located 247 bp upstream from the putative start codon.

### 3.4 Lineage-specific distribution of MntR miniregulon members

To assess the evolutionary conservation of the proposed MntR miniregulon, we analyzed the distribution of its components across 103 RefSeq genomes representing the diversity of *Stenotrophomonas* and related *Lysobacteraceae* (Fig. 4). The presence-absence patterns of MntR, MntH, MntP, ORF2357, and ORF2358, together with the occurrence of predicted MntR-binding motifs, revealed a non-uniform distribution across the analyzed taxa. Notably, the complete five-gene set, along with three associated MntR-binding sites, was exclusively detected in *Stenotrophomonas* species, including *S. maltophilia*. In contrast, closely related species such as *S. pavanii*,*S. beteli*, and *S. hibiscicola* lacked the TBDR and pTFP homologs and contained only two predicted MntR-binding motifs (Fig. 4). Furthermore, these genes were absent from the genomes of *Allostenotrophomonas*, *Pseudostenotrophomonas*, and *Parastenotrophomonas* (Chauviat et al., 2025). Among the components analyzed, only the MntP exporter was conserved across all four genera. Mapping these patterns onto the species phylogeny revealed that the complete (three-loci) MntR miniregulon is restricted to the *S. maltophilia* complex (Fig. 4). Importantly, the consistent co-occurrence of these genes with conserved MntR-binding motifs supports the existence of a lineage-specific, coordinately regulated gene module. These findings prompted us to experimentally validate the metal-dependent regulation of these loci.

**Fig. 4.**
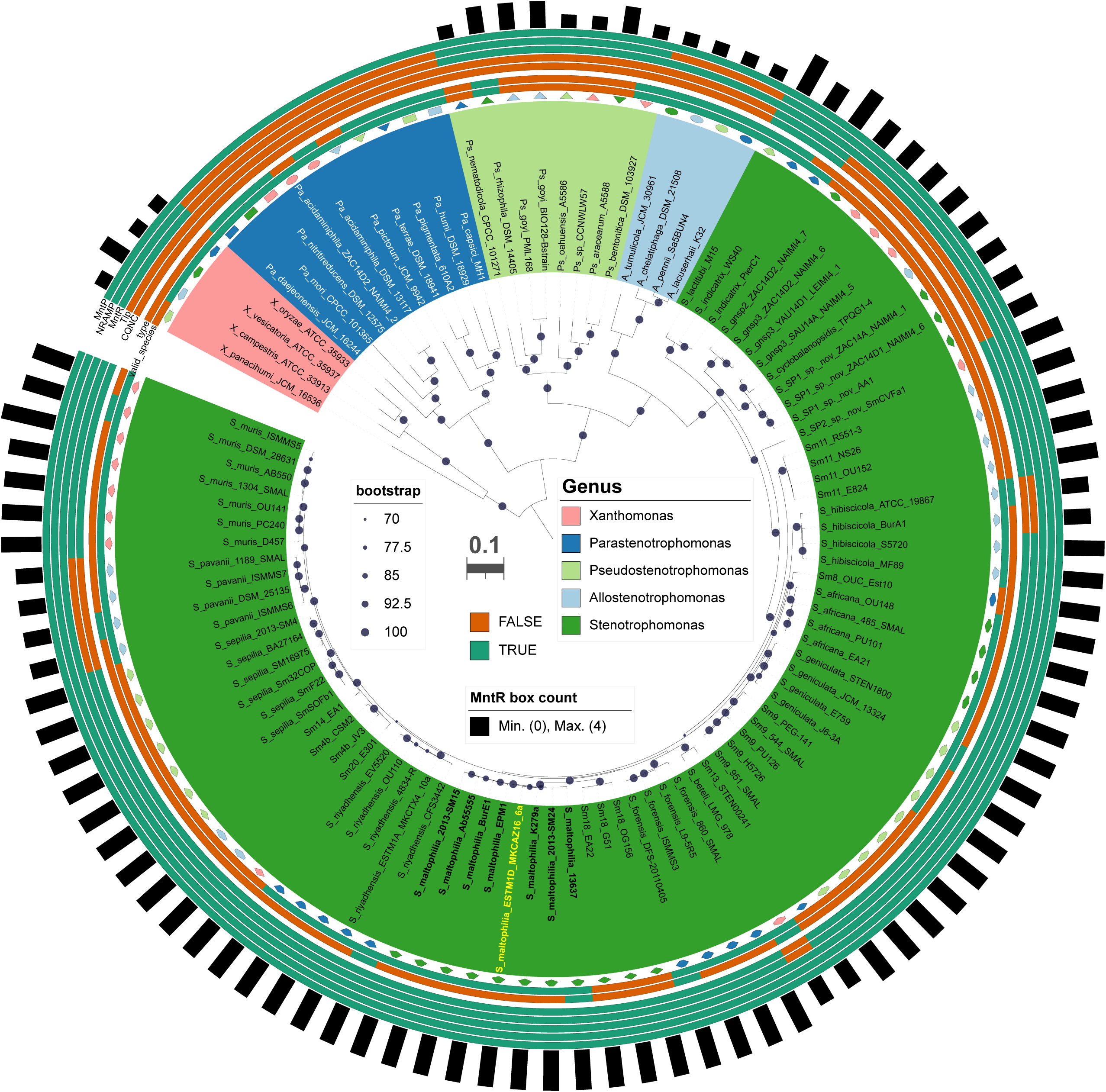
Distribution of the MntR miniregulon proteins across *Stenotrophomonas* species and closely related genera in the Lysobacteraceae family. Maximum-likelihood core-genome phylogeny estimated with IQTree based on the concatenation of 155 top-scoring protein families selected with GET_PHYLOMARKERS from 103 Lysobacteraceae RefSeq genomes (see color code). The tree integrates taxonomic information (four inner rings) with the presence/absence patterns of the MntR miniregulon genes (*orf02357*, *orf02358, mntH*, *mntR*, *and mntP*) identified by hmmsearch (five outer rings), and the number of MntR-boxes detected by MAST scanning (bars). *S. maltophilia* strains are labeled in bold font, with Sm18 highlighted in yellow. Bipartition support values are given as UFBoot bootstrap percentages only for those ≥ 70%. The scale bar represents the number of expected substitutions per site under the best-fitting Q.insect+F+R4 model, as selected by BIC.

### 3.5 The MntR miniregulon expression is modulated by iron and manganese

To validate the metal-dependent regulation of MntR miniregulon elements, we constructed transcriptional fusions of the 5’-UTRs of *orf2357-orf2358*, *mntH,* and *mntP* to evaluate their expression in the Sm18 background. Expression data for the Sm18/327-2357_2358 fusion revealed that it was exclusively expressed under simultaneous Mn and Fe limitation, with reporter activity completely abolished by a low Mn^2+^ concentration (80 nM) (Fig. 5A, first panel). Activity of the Sm18/327-*mntH* fusion was negatively correlated with increasing Mn^2+^ concentrations in the culture medium. However, low activity was still measurable at 8 µM Mn^2+^, and a significantly higher expression was observed at Fe^2+^ 10 µM compared with Fe^2+^ 0 µM, except when the medium contained 8 µM Mn^2+^ (Fig. 5A, second panel). Finally, Sm18/327-*mntP* increased significantly at 8 µM Mn^2+^ compared to Mn^2+^ 0µM, irrespective of Fe availability in the medium (Fig. 5A, third panel). Overall, these results show that the expression of *orf2357-orf2358*, *mntH,* and *mntP* is differentially modulated by Mn^2+^ and Fe^2+^ availability, with distinct effects on each locus. Notably, the transcriptional responses observed in these reporter assays are consistent with the trends identified in the RNA-seq analysis, validating the described regulatory effects of both transition metals on the MntR miniregulon members.

**Fig. 5.**
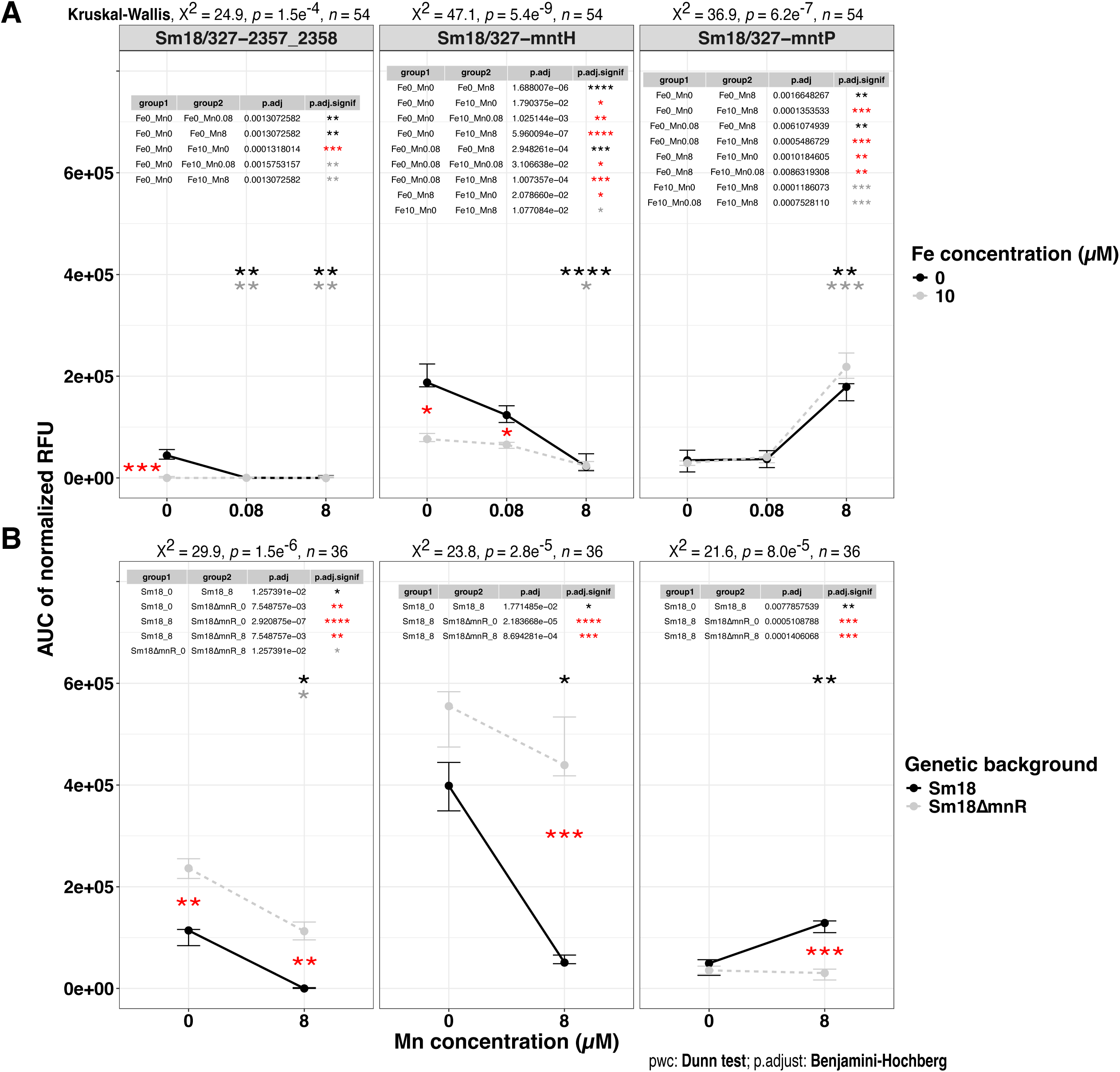
Statistical analysis of fluorescence intensity for transcriptional fusions of Sm18 genes within the manganese MntR miniregulon. (A) Area under the curve (AUC) of normalized relative fluorescence units (RFU), normalized using the RFU of strains carrying the empty pSEVA-327 plasmid. Nine biological replicates were grown in MM supplemented with 30 µg/mL of chloramphenicol. AUC values of transcriptional fusions Sm18/327-2357_2358, Sm18/327-*mntH*, and Sm18/327-*mntP* are shown as mean ± 95% CI (bootstrap) for 0 and 10 µM Fe^2+^. Each Fe condition was tested with 0, 0.08, and 8 µM of Mn^2+^. (B) AUC of normalized RFU for five biological and four technical replicates of Sm18 and Sm18ΔmntR strains harboring the same fusions grown in MM-Fe0. Differences were assessed using the Kruskal-Wallis test followed by Dunn’s *post hoc* test with BH correction. Significance levels are shown as: ns, *p* > 0.05; *, *p* < 0.05; **, *p* < 0.01; ***, *p* < 0.001; ***, *p* < 0.0001. Black asterisks denote comparisons across Mn concentrations, red asterisks denote Fe comparisons, and gray asterisks indicate combined Fe/Mn variations.

### 3.6 MntR controls core manganese homeostasis genes in Sm18

To confirm that the promoter activity observed in the transcriptional fusions reported in Fig. 5A is regulated by MntR, and to further validate the core MntR miniregulon, the three fusions were transferred into a Sm18*ΔmntR* strain, and promoter activity was measured in MM-Fe0Mn0 and MM-Fe0Mn8 medium. The Sm18Δ*mntR*/327-2357_2358 fusion revealed a significant increase in expression compared with Sm18/327-2357_2358. However, adding manganese led to reduced expression, following a similar trend to that observed in the Sm18 background, although not to the same extent (Fig. 5B, first panel). Similarly, derepression of the 327-*mntH* fusion was observed in the Sm18*ΔmntR* genetic background compared to the wild-type. However, no statistically significant activity was detected upon addition of 8 µM Mn^2+^ (Fig. 5B, second panel). Finally, no expression was detected for Sm18Δ*mntR*/327-*mntP* independent of manganese concentration (Fig. 5B, third panel). Overall, these results indicate that the expression of *orf2357–orf2358, mntH,* and *mntP* is dependent on MntR, with distinct expression patterns observed for each locus in the Δ*mntR* background. Notably, *orf2357–orf2358* and *mntH* show increased expression in the absence of MntR, whereas *mntP* expression is not detected under these conditions.

### 3.7 The orf2357-orf2358 locus encodes a novel protein module associated with manganese homeostasis

Following the identification of *orf2357* and *orf2358* as members of the MntR miniregulon, we performed a structural and phylogenomic characterization of their encoded proteins. *Orf2357* encodes a TonB-dependent receptor (TBDR) with a predicted signal peptide and a canonical β-barrel architecture containing an N-terminal plug domain (Fig 6A). Notably, this domain harbors a conserved CQNC motif (Fig. 6B y C). Homology searches revealed a restricted phylogenetic distribution of closely related sequences, which cluster into a well-supported lineage characterized by the presence of the CQNC motif. These proteins are sparsely distributed across diverse bacterial taxa. Structural modeling strongly suggests that this motif, together with residues from the β-barrel domain, may coordinate a Mn^2+^ ion in a tetrahedral geometry (Fig. S2). The downstream gene, *orf2358,* encodes a predicted periplasmic thioredoxin-fold protein (pTFP) (Nilewski et al., 2021). Phylogenomic analysis revealed that pTFP homologs form a distinct and well-supported clade with a limited taxonomic distribution (Fig. S3). Notably, these homologs are consistently found in synteny with CQNC-type TBDRs, reinforcing their functional linkage (Fig 6D). Sequence analysis of pTFP identified a conserved thioredoxin fold with a non-canonical CXXC motif (CAPC), embedded within a distinctive lineage-specific sequence signature (Fig. 6E, S3).Together, these findings identify a novel, MntR-regulated protein module comprising a CQNC-type TBDR and an associated thioredoxin-fold protein, which are consistently found in synteny, with a restricted phylogenetic distribution.

**Fig. 6.**
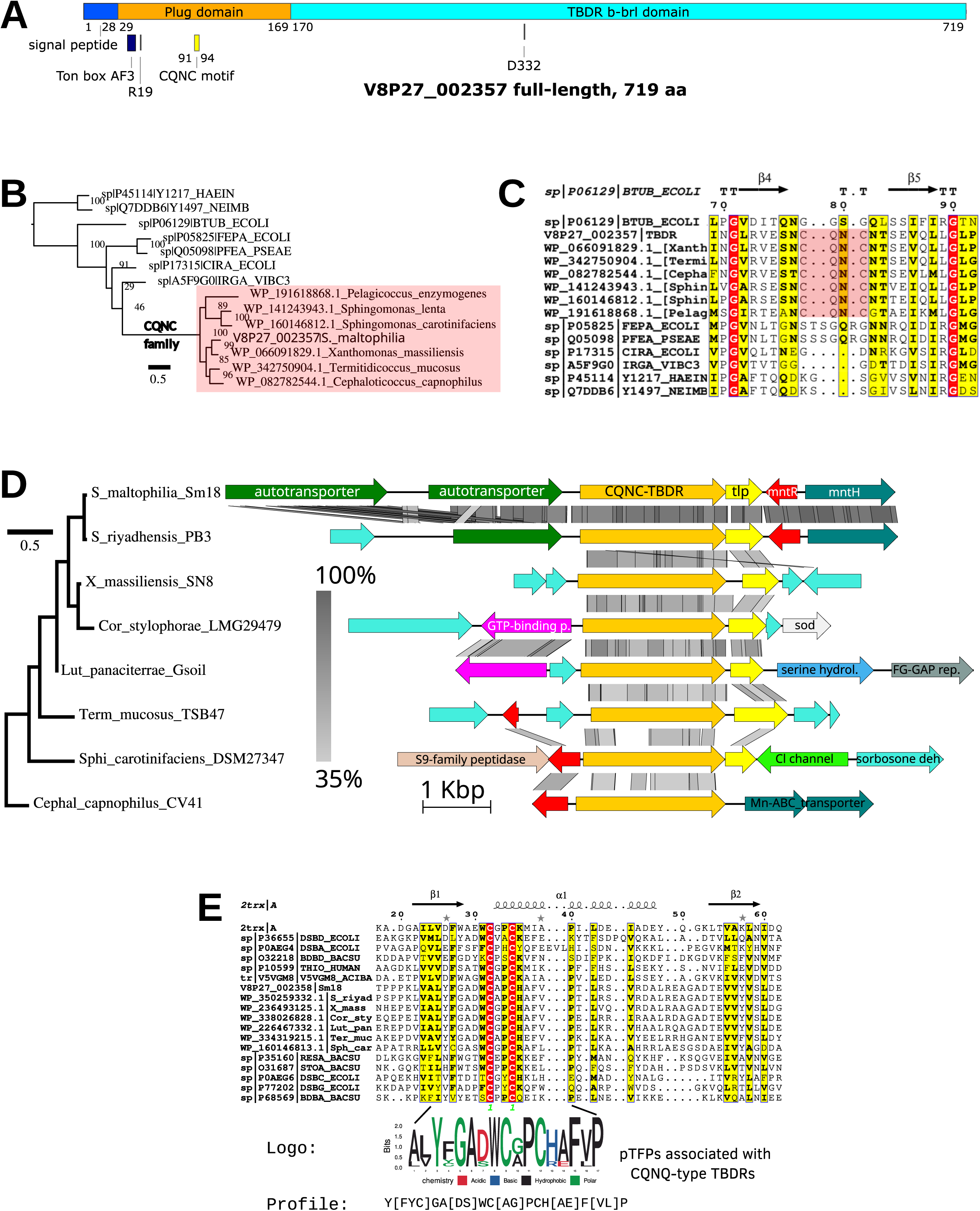
Structural, phylogenomic and comparative analysis of the novel CQNC family of TBDRs and associated periplasmic thioredoxin-fold protein (pTFP). (A) Domain organization of V8P27_002357 (TBDR) showing the β-barrel and plug domains, the N-terminal signal peptide and the location of the CQNC motif. (B) Maximum-likelihood phylogeny showing the monophyly and restricted taxonomic distribution of the CQNC-TBDR. (C) ESPript alignment showing the conserved CQNC motif located in the center of the plug domain. (D) Comparative genomics of the TBDR-pTFP showing conserved synteny of TBDR-pTFP loci across Lysobacterales, Sphingomonadales, and Opitutales. Gray shading indicates sequence identity. Loci are ordered according to a neighbor-joining tree based on Mash distances. E) Structure-guided multiple-sequence alignment and a close-up of the CXXC motif of the pTFP (V8P27_002358), with logo and profile (in PROSITE format) of the presumed catalytic cysteines (C39, C42) and neighboring residues.

### 3.8 MntR miniregulon mutants display sensitivity to hydrogen peroxide and manganese

To determine whether the *mntH* and *mntP* homologs in Sm18 encode functional manganese importer and exporter proteins, respectively, we constructed mutants of both genes and evaluated their growth under different stress conditions in MM-Fe0Mn0. The growth kinetics of the Sm18VIM*mntH* mutant indicated that the *mntH* gene is not essential under Fe^2+^- or Mn^2+^-limiting conditions in axenic cultures. However, it is required for growth under oxidative stress induced by hydrogen peroxide (H_2_O_2_), particularly at neutral pH (Fig. 7A). Statistical analysis revealed that the difference in growth was significant only at 500 µM H_2_O_2_ compared to the wild type (Fig. 7B). Strain Sm18Δ*mntP* was challenged with increasing Mn^2+^ concentrations (5-160 µM; Fig. 7C), with statistically significant sensitivity detectable at 40 µM, and complete growth inhibition observed at 160 µM (Fig. 7D). The wild-type phenotypes of the Sm18VIM-*mntH* and Sm18Δ*mntP* strains were restored by plasmids Sm18VIM-*mntH*/332::*mntH* and Sm18Δ*mntP*/332::*mntP* provided in *trans*, and carrying the *mntH* and *mntP* genes under the control of their native promoters (Fig. 7). These results show that *mntH* and *mntP* play key roles in resistance to H_2_O_2_ and manganese toxicity, respectively, as supported by the stress-sensitive phenotypes of the mutants and their restoration upon complementation. No growth phenotypes were detected for the markerless deletion *orf2357* and *orf2358* mutants under the conditions tested.

**Fig. 7.**
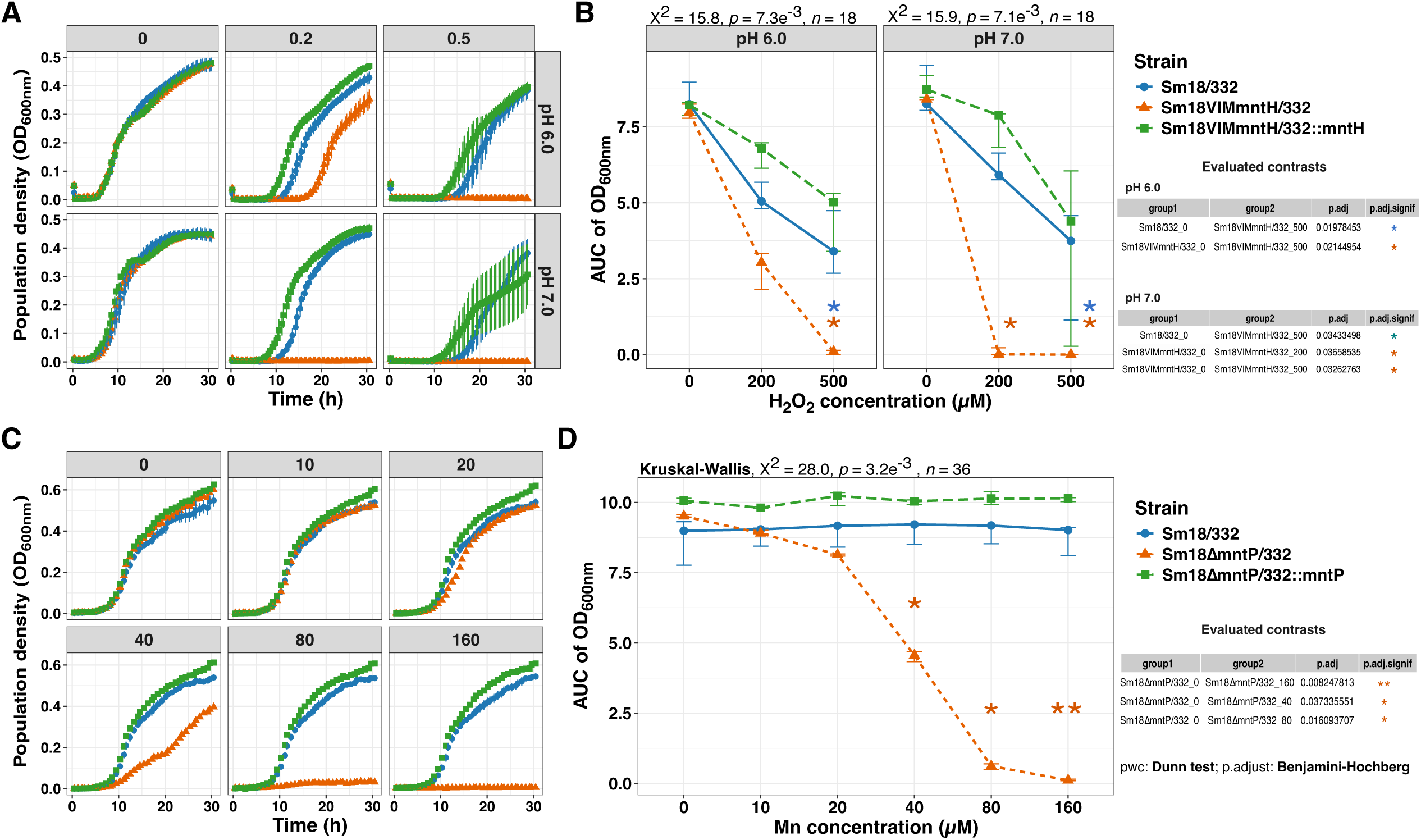
Phenotypic characterization of *S. maltophilia* Sm18 *mntH* and *mntP* mutants. (A-B) Growth kinetics of the Sm18VIM-*mntH* mutant in MM-MES under oxidative stress induced by hydrogen peroxide (H_2_O_2_) at pH 7.0 and 6.0. The *mntH* mutant (orange triangles) exhibited increased sensitivity to hydrogen peroxide (0.2 and 0.5 mM), particularly at neutral pH. Providing pSEVA332::*mntH* in *trans* restores growth to wild-type levels. (C-D) Growth kinetics of the markerless Sm18Δ*mntP* mutant (orange triangles) exposed to increasing Mn^2+^ concentrations (5-160 µM). Deletion of *mntP* results in significant manganese sensitivity starting at 40 µM, with complete growth inhibition at higher concentrations. Plasmid pSEVA332::*mntP* expressing *mntP* under its native promoters restored the wild type phenotype (green cubes). Error bars represent SEM for growth kinetics and 95% CI (bootstrap) for AUC analysis for three biological replicates. Statistical significance was evaluated using the Kruskal-Wallis test with Dunn’s *post hoc* BH correction: ns, *p* > 0.05; *, *p* < 0.05; **, *p* < 0.01; ***, *p* < 0.001; ****, *p* < 0.0001. Blue asterisks denote comparisons between wild type (blue circles) and mutants; orange asterisks indicate comparisons across H_2_O_2_ or Mn²⁺ concentrations.

### 3.9 The S. maltophilia mntH gene is expressed and supports intracellular replication in A. castellanii

We recently found that *S. maltophilia* Sm18 resists digestion by *A. castellanii* strain Neff trophozoites and can replicate in acidified, Rab7A-positive *Stenotrophomonas*-containing vacuoles (SCV) of this free-living professional phagocyte (Rivera et al., 2024). Given that the phagosomes of *Dictyostelium discoideum* and human macrophages subject bacteria to iron and manganese limitation via the NRAMP exporter (Morey et al., 2015; Peracino et al., 2013), we postulated that the Sm18 MntH Mn importer is required to survive in the SCV. To test this hypothesis, we first evaluated the activity of the Sm18/337-*mntH* fusion in *A. castellanii* strain Neff trophozoites using wide-field fluorescence microscopy. As shown in Fig. 8A, the fusion was expressed at 3 and 18 h post-primary contact (ppc). In contrast, equivalent assays performed with the Sm18/337R-*2357_2358* and Sm18/337R-*mntP* fusions did not reveal detectable expression under the same conditions. Moreover, the replication capacity of Sm18 in trophozoites was compared with that of the Sm18VIM*mntH* (*VIMmntH*) and Sm18Δ*mntR* (*ΔmntR*) mutants. As shown in Fig. 8B, the *VIMmntH* mutant had a significantly reduced replication capacity in Neff trophozoites, whereas that of *ΔmntR* was indistinguishable from Sm18 (Figs. 8B and 8C). Importantly, no differences in growth were observed among these strains under *in vitro* conditions in MM or LB (Fig. S5). Together, these observations indicate that the replication defect of the *VIMmntH* mutant is specific to the intracellular environment.

**Fig. 8.**
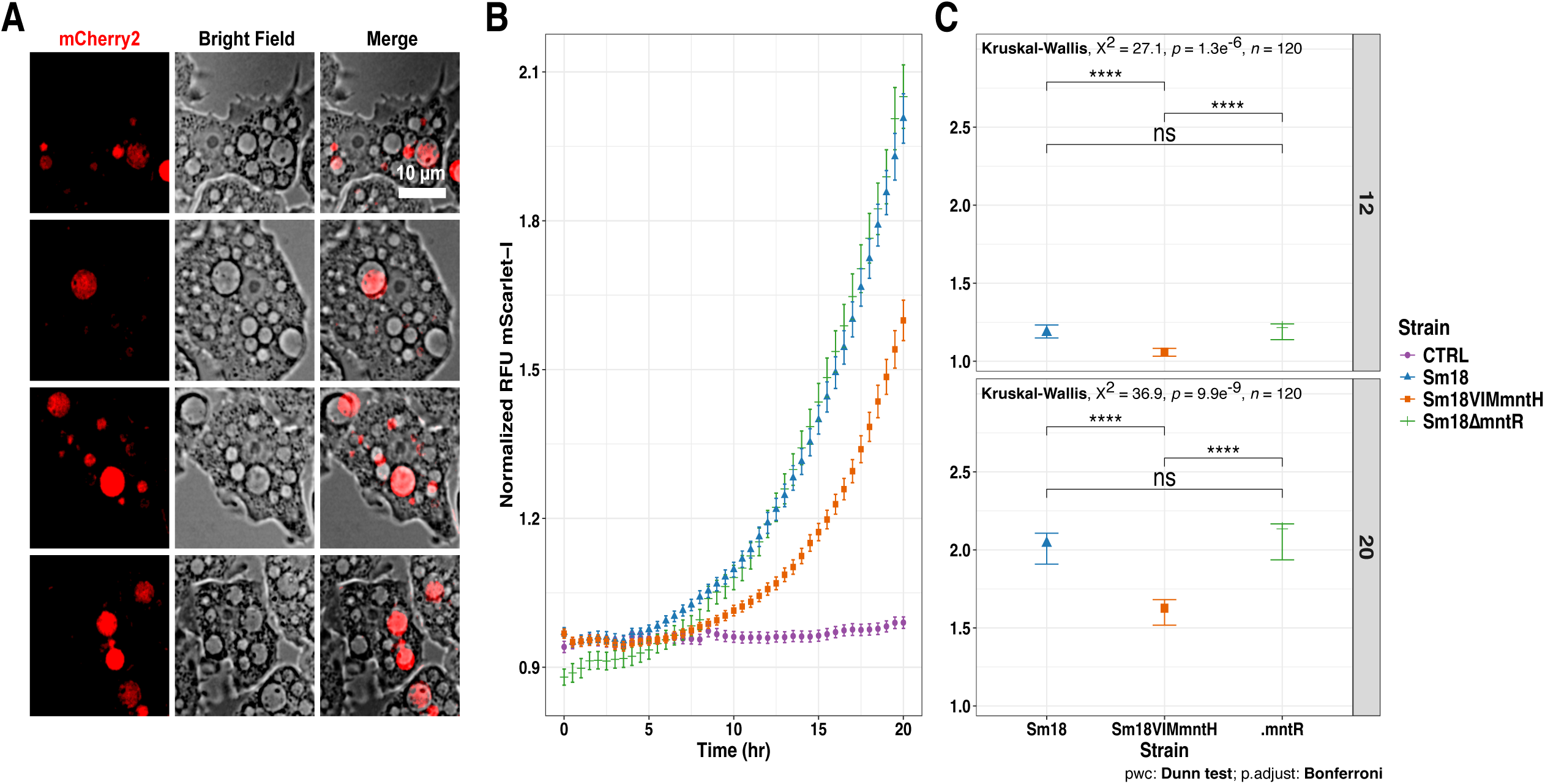
Live-cell imaging of Sm18/337-*mntH* promoter activity and intracellular replication of the Sm18VIM-*mntH* mutant in co-culture with *A. castellanii* trophozoites. (A) Representative images from six biological replicates of Sm18/337R-*mntH* and *A. castellanii* co-cultures at 3 and 18 h post primary contact. Promoter activity was visualized in the red channel (mCherry2). Trophozoites were imaged using brightfield microscopy. (B) Fluorescence kinetics of mScarlet-I-labeled strains: Sm18 (blue triangles), Sm18VIM*mntH* (orange cubes), and Sm18Δ*mntR* (green lines). Background fluorescence from medium-bacteria controls was subtracted for normalization. (C) Kruskal-Wallis test was used to compare medians between groups. When significant differences were detected, pairwise comparisons were performed using Dunn’s test with a Bonferroni correction. Significance levels: ns, *p* > 0.05; *, *p* < 0.05; **, *p* < 0.01; ***, *p* < 0.001; ****, *p* < 0.0001.

## 4. Discussion

The ability to maintain manganese homeostasis is critical for bacterial survival under fluctuating environmental and host-imposed conditions. In this study, we identified and characterized a regulatory and functional network that supports manganese homeostasis in *S. maltophilia* Sm18, centered on the metalloregulator MntR and a set of core target genes. Our results demonstrate that MntR plays a central role in modulating the expression of key manganese homeostasis genes, including *mntH*, *mntP*, and the novel *orf2357-orf2358* locus. Transcriptional analyses in the *ΔmntR* background revealed a clear derepression of *mntH* and the *orf2357-orf2358* putative operon in the absence of manganese, supporting a model in which MntR acts predominantly as a repressor under Mn-replete conditions. However, the persistence of manganese-responsive expression patterns in the *ΔmntR* strain suggests that additional regulatory layers contribute to fine-tuning this system. Thus, while our data establish MntR as a major regulator, they also indicate that manganese homeostasis in Sm18 is controlled by a more complex regulatory network that remains to be fully elucidated.

Beyond regulation, our functional analyses provide direct evidence for the physiological roles of MntR-regulated genes. The *VIMmntH* mutant displayed increased sensitivity to oxidative stress, consistent with the established role of manganese importers in protecting against reactive oxygen species (ROS). In contrast, deletion of *mntP* resulted in marked sensitivity even at low manganese concentrations, confirming its role in manganese detoxification. The restoration of wild-type phenotypes upon genetic complementation further supports the specific involvement of these genes in manganese homeostasis. Together, these findings validate the functional relevance of the MntR-controlled gene set and support the existence of a coherent regulatory module governing manganese uptake and efflux.

In addition to these canonical components, we identified the *orf2357-orf2358* locus as a previously uncharacterized element associated with manganese homeostasis. Structural and phylogenomic analyses suggest that *orf2357* encodes a TonB-dependent receptor with a conserved CQNC motif, while *orf2358* encodes a thioredoxin-fold protein that is consistently found in synteny with CQNC-type receptors, and might be function as a specialized disulfide reductase of the CQNC motif. The restricted phylogenetic distribution and conserved genomic association of these proteins support the existence of a distinct protein module. Although structural modeling suggests a potential role in manganese coordination, the precise function of this system remains to be experimentally validated. Therefore, our findings provide a foundation for future studies aimed at elucidating the mechanistic contribution of this module to manganese homeostasis.

Importantly, our results extend the relevance of manganese homeostasis to a biologically meaningful context. We show that *mntH* is expressed during interaction with *A. castellanii* and is required for optimal intracellular replication. The impaired replication of the *VIMmntH* mutant in amoebal trophozoites, together with its normal growth under *in vitro* conditions, indicates that this phenotype is specifically linked to the intracellular environment. These observations are consistent with previous reports showing that phagosomal compartments impose metal limitation as an antimicrobial strategy.

In parallel, the increased sensitivity of the *VIMmntH* mutant to H_2_O_2_ observed *in vitro* indicates that manganese uptake via MntH contributes to oxidative stress resistance. Although oxidative stress was not directly measured during amoeba infection, these findings support a model in which MntH-mediated manganese import enhances the bacterial capacity to cope with host-derived ROS within the phagosomal environment. In this context, manganese likely acts as a cofactor for antioxidant systems, including catalases and other ROS-detoxifying enzymes, thereby sustaining bacterial survival under oxidative stress conditions.

Consequently, impaired manganese acquisition in the *VIMmntH* mutant is expected to compromise these protective mechanisms, leading to reduced intracellular replication. This integrated model linking manganese homeostasis, oxidative stress defense, and intracellular survival is summarized in Fig. 9.

**Fig. 9.**
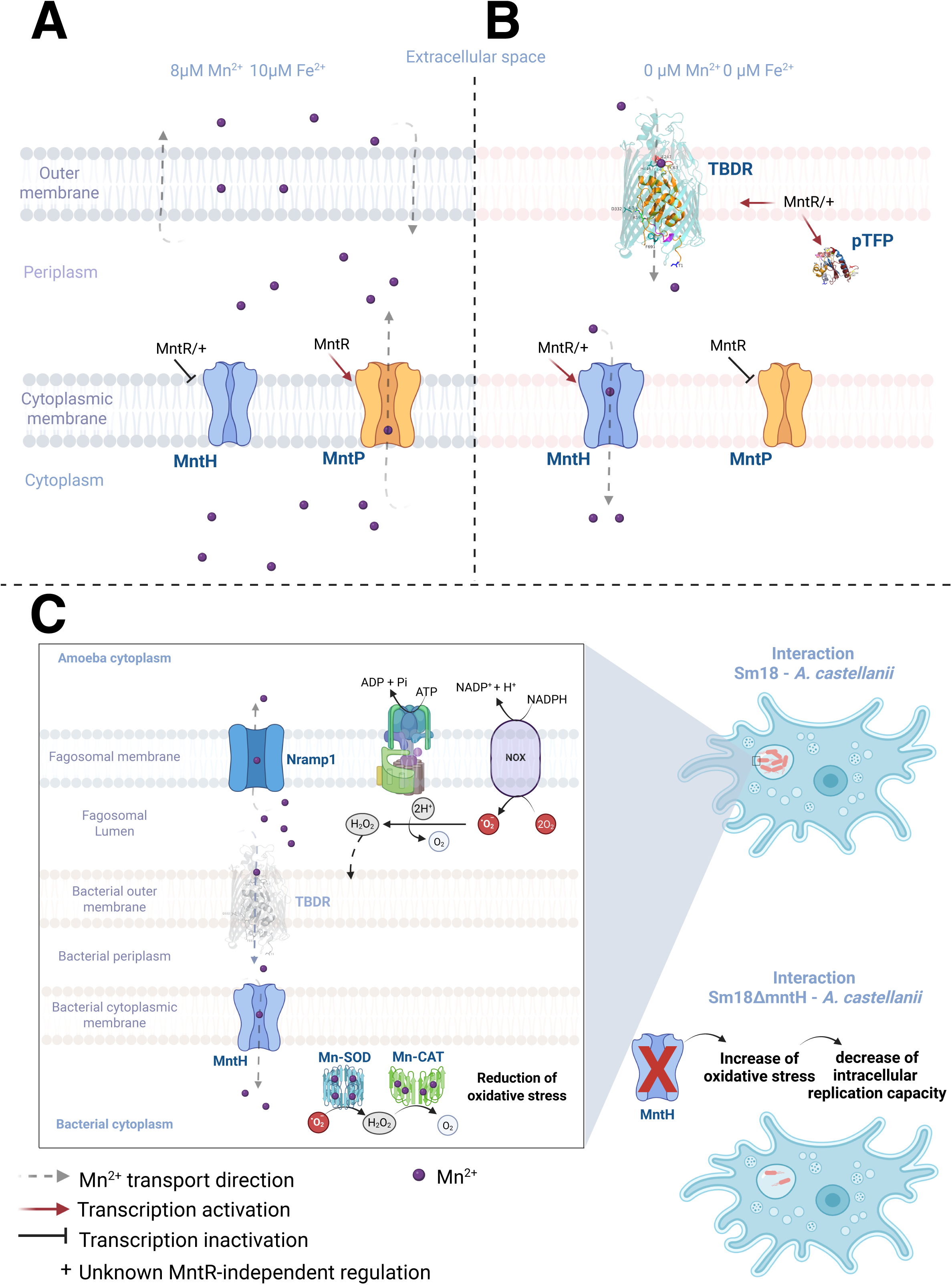
Model of *S. maltophilia* manganese homeostasis and its role during the interaction with *A. castellanii* trophozoites. (A) Schematic representation of manganese transport and gene regulation in strain Sm18 under defined *in vitro* conditions with varying Mn^2+^ and Fe^2+^ concentrations. MntH and MntP mediate manganese uptake and efflux, respectively. (B) Genes associated with manganese homeostasis that are specifically induced under combined Mn and Fe limitation. Gene expression is regulated by MntR, with evidence from this study supporting the existence of an additional MntR-independent regulatory mechanism. (C) Model of Sm18 interaction with *A. castellanii*. Within the phagosomal environment, manganese uptake via MntH contributes to resistance against oxidative stress generated by host-derived H_2_O_2_. Disruption of *mntH* results in reduced intracellular replication. TBDR is shown with reduced opacity to indicate lack of detectable expression under these conditions. Created with https://BioRender.com.

Taken together, this study defines a manganese homeostasis network in *S. maltophilia* Sm18 that integrates transcriptional regulation by MntR, functional roles of manganese transport systems, and a previously unrecognized protein module. While our findings establish the importance of this system under both environmental and intracellular conditions, they also highlight the existence of additional regulatory and functional components that warrant further investigation.

## 5. Conclusion

In this study, we defined a manganese homeostasis network in *S. maltophilia* Sm18 centered on the metalloregulator MntR and its target genes. Our results demonstrate that *mntH* and *mntP* play key roles in oxidative stress resistance and manganese detoxification, respectively, while also identifying a previously uncharacterized TBDR-pTFP module associated with this regulatory system. Importantly, we show that manganese acquisition via MntH is required for optimal intracellular replication in *A. castellanii*, linking metal homeostasis with bacterial adaptation to host-associated environments. Overall, this work established a functional connection between manganese homeostasis, oxidative stress defense, and intracellular survival, providing a framework for understanding how *S. maltophilia* adapts to the metal-restricted and oxidative environment of phagosomes.

## CRediT authorship contribution statement

**Fulvia Stefany Argueta Zepeda:** Writing – original draft, Writing – review & editing, Validation, Methodology, Investigation, Formal analysis, Data curation. **Javier Rivera Campos:** Writing – review & editing, Validation, Methodology. **Julio Valerdi Negreros:** Writing – review & editing, Validation, Methodology. **Christopher Rensing:** Writing – review & editing, Supervision. **Pablo Vinuesa:** Writing – review & editing, Writing – original draft, Supervision, Resources, Project administration, Methodology, Investigation, Funding acquisition, Formal analysis, Conceptualization

## Conflict of interest

The authors declare no conflict of interest.

## ACKNOWLEDGEMENTS

We thank Alfredo Hernández and Víctor del Moral for their expert technical support and server maintenance at the Centro de Ciencias Genómicas, UNAM. We acknowledge the excellent service provided by personnel at the Unidad de Síntesis y Secuenciación de DNA from Instituto de Biotecnología, UNAM. We thank Dr. Ayari Fuentes from CCG-UNAM for providing access to the multimodal plate reader. We thank Dr. Luis Lozano from the Bioinformatics Unit at CCG-UNAM for his advice and support during the early phase of transcriptome analysis. We thank M.Sc. Laura Cervantes for her laboratory assistance and valuable feedback. We gratefully acknowledge the financial support received from Consejo Nacional de Humanidades Ciencias y Tecnologías (CONAHCyT México, A1-S-11242), Secretaría de Ciencia, Humanidades, Tecnología e Innovación (SECIHTI México CBF-2025-I-1106), and Programa de Apoyo a Proyectos de Investigación e Innovación Tecnológica (PAPIIT), Universidad Nacional Autónoma de México (DGAPA-PAPIIT, UNAM: IN209321 and IN216424) to PV. F. S. A. Z. and J.C.V.N. were recipients of CONAHCyT scholarships 659308 and 745705, respectively.

## Appendix A. Supporting information

Supplementary data associated with this article can be found in the online version at doi:10.1016/j.micres.2026 ….

## Data availability

The complete genome sequence of *S. maltophilia* Sm18 was deposited in GenBank under accession number CP146374 and the associated genome and transcriptome sequencing reads were deposited under BioProject PRJNA1081934 https://www.ncbi.nlm.nih.gov/bioproject/?term=PRJNA1081934. The RefSeq genome assembly for strain Sm18 is publicly available from NCBI’s genome datasets portal at https://www.ncbi.nlm.nih.gov/datasets/genome/GCA_053078355.1/.

The sequence reads generated from the 18 RNAseq samples analyzed in this work were deposited at NCBI’s Sequence Read Archive https://www.ncbi.nlm.nih.gov/sra/ under accession numbers SRR36898890 to SRR36898907, as part of BioProject PRJNA1081934 https://www.ncbi.nlm.nih.gov/bioproject/?term=PRJNA1081934.

The hidden Markov models built in this work are freely available from GitHub at: https://github.com/vinuesa/supplementary_materials/tree/main/docs/Argueta-Zepeda_Stenotrophomonas_MntR_miniregulon.

## Conflict of interest disclosure statement

None declared.

## Ethics statement

None required.

## Notes

https://github.com/vinuesa/supplementary_materials/blob/main/docs/Argueta-Zepeda_Stenotrophomonas_MntR_miniregulon/Argueta-Zepeda_suppl_tables.pdf

https://www.ncbi.nlm.nih.gov/bioproject/1081934

### Competing Interest Statement

The authors have declared no competing interest.

### Summary of Updates

The title has been revised to more accurately reflect the study's scope and to align with the version currently under peer review. Significant edits were performed throughout the manuscript to make it more succinct. The reference for reporting the complete genome sequence for strain Sm18 was updated. Figure 9 was also updated.

