## Supplemental tables S1-S6 for "Manganese Homeostasis Drives *Stenotrophomonas maltophilia* Oxidative Stress Defense and Replication in *Acanthamoeba castellanii* phagosomes"

**Table S1.** Sm18 significantly up- and down-regulated genes from 0 µM to 8 µM of Mn(II) in minimal MOPS medium with 10 µM Iron. Cutoff: adjusted p-value (padj) < 0.01 and Log_2_ fold change (Log_2_FC) ≥ 1 or ≤ -1.

| **Locus id** | **Gene name** | **Log_2_FC** | **DE** | **padj** | **Gene description** |
| --- | --- | --- | --- | --- | --- |
| V8P27_004008 | *mntP* | 3.4 | Up | 4.2 e^-61^ | Manganese efflux pump |
| V8P27_003797 | V8P27_003797 | 2.0 | Up | 1.4 e^-08^ | Copper chaperone SCO1/SenC |
| V8P27_003805 | V8P27_003805 | 1.8 | UP | 1.4 e^-08^ | HTH-type transcriptional regulator |
| V8P27_003804 | V8P27_003804 | 1.7 | UP | 6.0 e^-03^ | DUF6436 domain-containing protein |
| V8P27_002360 | *mntH* | 1.0 | Down | 2.0 e^-05^ | Mn2+/Fe2+ uptake protein |

**Table S2.** Sm18 significantly up- and down-regulated genes from 0 µM to 8 µM of Mn(II) in minimal MOPS medium without Iron. Cutoff: adjusted p-value (padj) < 0.01 and Log_2_ fold change (Log_2_FC) ≥ 1 or ≤ -1.

| **Locus id** | **Gene name** | **Log_2_FC** | **DE** | **padj** | **Gene description** |
| --- | --- | --- | --- | --- | --- |
| V8P27_004008 | *mntP* | 4.0 | Up | 2.2 e^-131^ | Manganese efflux pump |
| V8P27_001285 | V8P27_001285 | 2.2 | Up | 4.0 e^-04^ | TonB-dependent receptor |
| V8P27_004004 | V8P27_004004 | 1.3 | Up | 1.8 e^-20^ | Major Facilitator Superfamily transporter |
| V8P27_004005 | V8P27_004005 | 1.3 | Up | 4.8 e^-11^ | DcaP outer membrane protein |
| V8P27_001286 | V8P27_001286 | 1.2 | Up | 3.7 e^-04^ | MerC domain-containing protein |
| V8P27_000822 | *prpE* | 1.0 | Up | 1.3 e^-09^ | propionate-CoA ligase |
| V8P27_002357 | V8P27_002357 | 4.6 | Down | 8.5e^-122^ | TonB-dependent receptor |
| V8P27_002358 | V8P27_002358 | 2.8 | Down | 5.3 e^-03^ | Thioredoxin-fold protein |
| V8P27_003990 | V8P27_003990 | 2.0 | Down | 1.0 e^-05^ | MFP of RND efflux pump |
| V8P27_001267 | V8P27_001267 | 1.8 | Down | 6.5 e^-03^ | Cation Diffusion Facilitator transporter |
| V8P27_002373 | V8P27_002373 | 1.5 | Down | 8.4 e^-09^ | Small Multidrug Resistance transporter |
| V8P27_003991 | V8P27_003991 | 1.5 | Down | 1.8 e^-14^ | RND efflux pump |
| V8P27_002360 | *mntH* | 1.5 | Down | 8.2 e^-15^ | Mn2+/Fe2+ uptake protein |
| V8P27_003989 | V8P27_003989 | 1.4 | Down | 4.0 e^-04^ | OMF of RND efflux pump |
| V8P27_001268 | *nfi* | 1.3 | Down | 3.6 e^-08^ | Endonuclease V |
| V8P27_001249 | V8P27_001249 | 1.3 | Down | 7.1 e^-07^ | Uncharacterized protein |
| V8P27_001248 | *map* | 1.2 | Down | 1.7 e^-12^ | Type I methionyl aminopeptidase |

**Table S3.** Strains used and constructed in this study.

| **Strain name** | **Key strain** | **Reference/source** |
| --- | --- | --- |
| DH5α | Standard *Escherichia coli* cloning strain. F− endA1 glnV44 thi-1 recA1 relA1 gyrA96 deoR nupG purB20 | Laboratory strain collection |
| S17.1 | Standard *Escherichia coli* conjugative donor strain. recA, thiE1, pro-82, endA, hadR17. RP4-2(Km::Tn7, Tc::Mu1) | (1) |
| Sm18 | *Stenotrophomonas maltophilia* strain Sm18. Environmental isolate from Cuernavaca, Morelos, México. | (2) |
| Neff | *Acanthamoeba castellanii* strain Neff. Environmental isolate from the USA | ATCC 30010 |
| Sm18∆*02357* | Deletion mutant of the ORF 02357 from *Stenotrophomonas maltophilia* Strain Sm18 | This work |
| Sm18∆*02358* | Deletion mutant of the ORF 02358 from *Stenotrophomonas maltophilia* Strain Sm18 | This work |
| Sm18∆*mntP* | Deletion mutant of the ORF mntP from *Stenotrophomonas maltophilia* Strain Sm18 | This work |
| Sm18∆*mntR* | Deletion mutant of the ORF mntR from *Stenothophomonas maltophilia* Strain Sm18 | This work |
| Sm18/332 | *Stenotrophomonas maltophilia* strain Sm18 transformed with the empty vector pSEVA332 | This work |
| Sm18∆*mntP*/332 | Sm18∆*mntP* deletion mutant transformed with the empty vector pSEVA332 | This work |
| Sm18∆*mntP*/332::*mntP* | Sm18∆*mntP* deletion mutant transformed with the vector pSEVA332::*mntP* | This work |
| Sm18::mTn7TC1_Pc_mScarlet-I | *Stenotrophomonas maltophilia* strain Sm18 derivative tagged with mScarlet-I expressed from the strong constitutive Pc promoter. | (3) |
| Sm18∆*02357*::mTn7TC1_Pc_mScarlet-I | Derivative of the Sm18∆*02357* deletion mutant, tagged with mScarlet-I and expressed under the strong constitutive Pc promoter | This work |
| Sm18∆*02358*::mTn7TC1_Pc_mScarlet-I | Derivative of the Sm18∆*02358* deletion mutant, tagged with mScarlet-I and expressed under the strong constitutive Pc promoter | This work |
| Sm18∆*mntP*::mTn7TC1_Pc_mScarlet-I | Derivative of the Sm18∆*mntP* deletion mutant, tagged with mScarlet-I and expressed under the strong constitutive Pc promoter | This work |
| Sm18∆*mntR*::mTn7TC1_Pc_mScarlet-I | Derivative of the Sm18∆*mntR* deletion mutant, tagged with mScarlet-I and expressed under the strong constitutive Pc promoter | This work |
| Sm18VIM*02357* | Interrupted mutant of the ORF *02357* from *Stenotrophomonas maltophilia* Strain Sm18 | This work |
| Sm18VIM*mntH* | Interrupted mutant of the ORF mntH from *Stenotrophomonas maltophilia* Strain Sm18 | This work |
| Sm18VIM*mntP* | Interrupted mutant of the ORF mntP from *Stenotrophomonas maltophilia* Strain Sm18 | This work |
| Sm18VIM*mntH*::mTn7TC1_Pc_mScarlet-I | Derivative of the Sm18VIMmntH interrupted mutant, tagged with mScarlet-I and expressed under the strong constitutive Pc promoter | This work |
| Sm18VIM*mntH*/332 | Sm18VIMmntH interrupted mutant transformed with the empty vector pSEVA332 | This work |
| Sm18VIM*mntH*/332::mntH | Sm18VIMmntH interrupted mutant transformed with the vector pSEVA332::*mntH* | This work |
| Sm18/327 | *Stenotrophomonas maltophilia* strain Sm18 transformed with the empty expression vector pSEVA327 | This work |
| Sm18/327-pr_*02357*-*02358* | *Stenotrophomonas maltophilia* strain Sm18 transformed with the expression vector pSEVA327-pr_*02357-02358* | This work |
| Sm18/327-pr_*mntH* | *Stenotrophomonas maltophilia* strain Sm18 transformed with the expression vector pSEVA327-pr_*mntH* | This work |
| Sm18/327-pr_*mntP* | *Stenotrophomonas maltophilia* strain Sm18 transformed with the expression vector pSEVA327-pr_*mntP* | This work |
| Sm18/337R | *Stenotrophomonas maltophilia* strain Sm18 transformed with the empty expression vector pSEVA337R | This work |
| Sm18/337R-pr_*02357*-*02358* | *Stenotrophomonas maltophilia* strain Sm18 transformed with the expression vector pSEVA337R-pr_*02357*-*02358* | This work |
| Sm18/337R-pr_*mntH* | *Stenotrophomonas maltophilia* strain Sm18 transformed with the expression vector pSEVA337R-pr_*mntH* | This work |
| Sm18/337R-pr_*mntP* | *Stenotrophomonas maltophilia* strain Sm18 transformed with the expression vector pSEVA337R-pr_*mntP* | This work |
| Sm18∆*mntR*/327 | Derivative of the Sm18∆*mntR* deletion mutant transformed with the empty expression vector pSEVA327 | This work |
| Sm18∆*mntR*/327-pr_*02357*-*02358* | Derivative of the Sm18∆mntR deletion mutant transformed with the expression vector pSEVA327-pr_*02357*-*02358* | This work |
| Sm18∆*mntR*/327-pr_*mntH* | Derivative of the Sm18∆mntR deletion mutant transformed with the expression vector pSEVA327-pr_*mntH* | This work |
| Sm18∆*mntR*/327-pr_*mntP* | Derivative of the Sm18∆mntR deletion mutant transformed with the expression vector pSEVA327-pr_*mntP* | This work |

**Table S4:** Vectors used and constructed in this study

| **Plasmid name** | **Plasmid features** | **Reference/source** |
| --- | --- | --- |
| pUC18T_mTn7TC1_Pc_mScarlet-I | pUC18T_mTn7TC1_Pr_mScarlet-I derivative for chromosomal labeling of bacteria with constitutive mScarlet-I expression driven from the strong Pc promoter; 5,893 bp. | (3) |
| pTNS2 | R6K-based plasmid (ApR) encoding the TnsABCD Tn7 transposase expression genes; 9,615 bp. | AddGene #64968 |
| pEX18Tc | Mobilizable plasmid containing OriT, for site-targeted mutagenesis in Gram-negative bacteria; 6349 bp. | (4) |
| Sm18∆*02357*_pEX18Tc | pEX18Tc derivative with homologous DNA genetic fragments flanking the ORF *02357* of Sm18; 7424 bp. | This work |
| Sm18∆*02358*_pEX18Tc | pEX18Tc derivative with homologous DNA genetic fragments flanking the ORF *02358* of Sm18; 7343 bp. | This work |
| Sm18∆*mntP*_pEX18Tc | pEX18Tc derivative with homologous DNA genetic fragments flanking the ORF *mntP* of Sm18; 7375 bp. | This work |
| Sm18∆*mntR*_pEX18Tc | pEX18Tc derivative with homologous DNA genetic fragments flanking the ORF *mntR* of Sm18; 7340 bp. | This work |
| pEX18TcVIM_GFP_PEM7 | pEx18Tc derivative with expression of GFP under control of constitutive promoter PEM7 and deletion of the gen SacB; 6031 pb. | Laboratory constructions collection |
| Sm18VIM*02357*_pEX18Tc∆SacB7_PEM7 | pEX18Tc∆*SacB7*_PEM7 derivative with homologous fragment of the ORF *02357* of Sm18; 6404 bp. | This work |
| Sm18VIM*mntH*_pEX18Tc∆SacB7_PEM7 | pEX18Tc∆*SacB7*_PEM7 derivative with homologous fragment of the ORF *mntH* of Sm18; 6380 bp. | This work |
| Sm18VIM*mntP*_pEX18Tc∆SacB7_PEM7 | pEX18Tc∆*SacB7*_PEM7 derivative with homologous fragment of the ORF *mntP* of Sm18; 6360 bp. | This work |
| pSEVA332 | Empty vector (chloramphenicol resistance, ori pBBR1, cargo lacZα-pUC19); 3417 bp. | (5) |
| pSEVA332::*mntH* | pSEVA332 digested with PacI and SpeI to remplace the lacZα-pUC19 cargo with mntH gene; 4452 bp. | This work |
| pSEVA332::*mntP* | pSEVA332 digested with PacI and SpeI to remplace the lacZα-pUC19 with *mntP* gene; 4077 bp. | This work |
| pSEVA327 | Empty expression vector (chloramphenicol resistance, ori RK2, cargo GFP); 4408 bp. | (5) |
| pSEVA327-pr_*02357*-*02358* | pSEVA327 containing the promoter region of the ORF *02357-02358*; 4796 bp. | This work |
| pSEVA327-pr_*mntH* | pSEVA327 containing the promoter region of the ORF *mntH;* 4721 bp. | This work |
| pSEVA327-pr_*mntP* | pSEVA327 containing the promoter region of the ORF *mntP*; 4976 bp. | This work |
| pSEVA337R | Empty expression vector (chloramphenicol resistance, ori pBBR1, cargo mCherry); 3661 bp. | (5) |
| pSEVA337R-pr_*02357*-*02358* | pSEVA337R containing the promoter region of the ORF *02357-02358*; 4049 bp. | This work |
| pSEVA337R-pr_*mntH* | pSEVA337R containing the promoter region of the ORF *mntH*; 3974 bp. | This work |
| pSEVA337R-pr_*mntP* | pSEVA337R containing the promoter region of the ORF *mntP*; 4229 bp. | This work |

**Table S5:** Primers designed in this study for constructing vectors

| **Primer name** | **Primer sequence (5’ to 3’)** | **Restriction sites** | **Reference/source** |
| --- | --- | --- | --- |
| Sm18_D02361_Fragment1.F | cgttgtaaaacgacggccagtgccatgtttggccagagc | None | This work |
| Sm18_D02361_Fragment1.R | tcgcagtgcttcacatgaaggcgtcgtcct | None | This work |
| Sm18_D02361_Fragment2.F | gacgccttcatgtgaagcactgcgagcgc | None | This work |
| Sm18_D02361_Fragment2.R | tctagagtcgacctgcaggcatgcatttcgcgcgaccc | None | This work |
| Sm18_D02362_Fragment1.F | tgtaaaacgacggccagtgccaaaaccaacctcggctgg | None | This work |
| Sm18_D02362_Fragment1.R | cgcgaccctaccaacagcgctcgcagt | None | This work |
| Sm18_D02362_Fragment2.F | gagcgctgttggtagggtcgcgcgaa | None | This work |
| Sm18_D02362_Fragment2.R | agagtcgacctgcaggcatgcatcgctgaaaagcaccccctt | None | This work |
| Sm18DmntP_PCR1.FOR | cgttgtaaaacgacggccagtgccacgcttacgacctcaacgtct | None | This work |
| Sm18DmntP_PCR1.REV | cgcggaagcgttaaatgggggacatggacagc | None | This work |
| Sm18DmntP_PCR2.FOR | atgtcccccatttaacgcttccgcgattacgt | None | This work |
| Sm18DmntP_PCR2.REV | ggaaacagctatgaccatgattacgcgatgttctcgatcattctgccg | None | This work |
| Sm18_DmntR_Fragment1.F | tgtaaaacgacggccagtgccaaacagcgctcgctgcgaccga | None | This work |
| Sm18_DmntR_Fragment1.R | gcgcaggcgtgtagcggtggcgccgg | None | This work |
| Sm18_DmntR_Fragment2.F | cgccaccgctacacgcctgcgcctg | None | This work |
| Sm18_DmntR_Fragment2.R | agagtcgacctgcaggcatgcaatcacttcggccaggtcgcaggcgatgatcgc | None | This work |
| Sm18_VIM_02361.F | aaaaaGAATTCcacacccctcttccatcgg | EcoRI | This work |
| Sm18_VIM_02361.R | aaaAAGCTTcttgaccacttcgatgcgg | HindIII | This work |
| Sm18-VIM-mntH.F | aaaaAAGCTTctacatgatctcggtcggct | HindIII | This work |
| Sm18-VIM-mntH.R | aaaaGGATCCgaagatcaccatcagcagcg | BamHI | This work |
| Sm18-VIM-mntP.F1 | aaaaAAGCTTccccatttcgatcctcctga | HindIII | This work |
| Sm18-VIM-mntP.R1 | aaaaGGATCCcgccgatatgcacatccatg | BamHI | This work |
| Sm18_mntHcompl.f | aaaaTTAATTAAtagtcctccaccagctccatc | PacI | This work |
| Sm18_mntHcompl.R | aaaaACTAGTaaggcatccacgcatggcg | SpeI | This work |
| Sm18_mntPcompl.F1 | aaaaTTAATTAAgcttacgacctcaacgtc | PacI | This work |
| Sm18_mntPcompl.R | aaaaACTAGTatcgcgtgctgaccaagcc | SpeI | This work |
| Sm18_pr_02357.F | aaaaaGAATTCtaccgcaaccatggagcc | EcoRI | This work |
| Sm18_pr_02357.R | aaaAAGCTTctgactttcacctgcacgc | HindIII | This work |
| Sm18-pr-mntH-p3.f | atatGGATCCtagtcctccaccagctccatc | BamHI | This work |
| Sm18-pr-mntH-p3.R | atatAAGCTTaaccaccagtgacccttgtcg | HindIII | This work |
| Sm18-pr-mntP.F1 | atataGAATTCgcttacgacctcaacgtc | EcoRI | This work |
| Sm18-pr-mntP.R1 | atatGGATCCgatcaggaggatcgaaatg | BamHI | This work |

**TABLE S6:** Primers used and synthesized in this study to verify constructions or mutants.

| **Primer name** | **Primer sequence (5’ to 3’)** | **Mutant/Constructions** | **Reference/source** |
| --- | --- | --- | --- |
| F24 | cgccagggttttcccagtcacgac | Forward primer for confirming constructs or merodiploids strains in plasmids pEXTc18 and constructs in pEXTc18∆SacB7_PEM7 | (6) |
| R24 | agcggataacaatttcacacagga | Reverse primer for confirming constructs in plasmids pEXTc18, insertional mutations in pEXTc18∆SacB7_PEM7 and Forward primer for confirming transcriptional fusions in pSEVA327 and pSEVA 337R | (6) |
| pBBR1_VER.F | cggccatcgtccacatatcc | Forward primer for confirming constructs in plasmid pSEVA332 | This work |
| Sm18_D02361_VER.F | atgttctgatcccgctttcg | Forward primer for confirming deletion mutant Sm18∆02357 | This work |
| Sm18_D02361_VER.R | tgacgtacaccacttcggtatc | Reverse primer for confirming deletion mutant Sm18∆02357 | This work |
| Sm18_D02362_VER.F | gctcgtggtacgtcaacctg | Forward primer for confirming deletion mutant Sm18∆02358 | This work |
| Sm18_D02362_VER.R | gtcgagctgatctcggacct | Reverse primer for confirming deletion mutant Sm18∆02358 | This work |
| Sm18_DmntP_VER.F | gctatacaaaccagccctgc | Forward primer for confirming deletion mutant Sm18∆mntP | This work |
| Sm18_DmntP_VER.R | tggatcaggcgttggagaaa | Reverse primer for confirming deletion mutant Sm18∆mntP | This work |
| Sm18_DmntR_VER.F | agcctggatgagagcgagag | Forward primer for confirming deletion mutant Sm18∆mntR | This work |
| Sm18_DmntR_VER.R | ttcatggggccgaatatagc | Reverse primer for confirming deletion mutant Sm18∆mntR | This work |
| Sm18_M02362_VER.R | tccaggctgacgtacaccac | Reverse primer for confirming merodiploids Sm18::pEX18Tc-02358 | This work |
| Sm18_MmntR_VER.R | agaccgtggagcgattcct | Reverse primer for confirming merodiploids Sm18::pEX18Tc-mntR | This work |
| Steno_glmS_down.F  (Smal_glmS_down_1549F) | gacatgccggtggtggtgatcg | Forward primer for confirming insertion of pUC18T_mTn7TC1_Pc_mScarlet-I in Sm18 Strains | (3) |
| pTn7.R | cacagcataactggactgatttc | Reverse primer for confirming insertion of pUC18T_mTn7TC1_Pc_mScarlet-I in Sm18 Strains | (3) |
