## Supplemental figures S1-S3 for "Manganese Homeostasis Drives *Stenotrophomonas maltophilia* Oxidative Stress Defense and Replication in *Acanthamoeba castellanii* phagosomes"

### Supplementary Figures S1-S3.

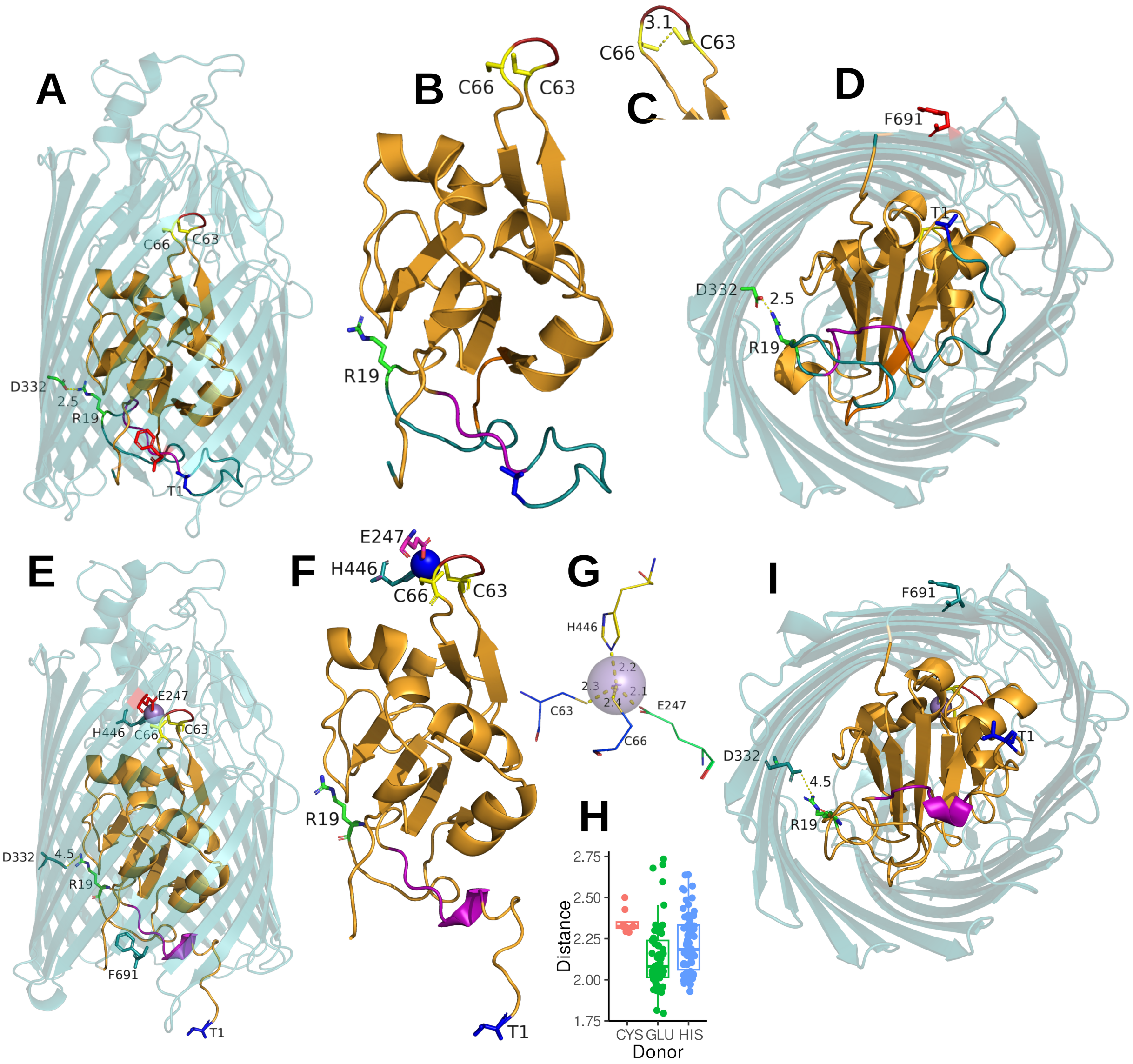


**Figure S1.** AlphaFold3 (Abramson et al., 2024) models of the *Stenotrophomonas maltophilia* V8P27_002357 protein encoding a novel family of TonB-dependent receptors (TBDRs) characterized by a CQNC motif in their plug domain, rendered with PyMOL (DeLano, 2004). (A-D) AF3 models of V8P27_002357 in its apo configuration (pTM = 0.93 and mean pLDDT = 88.4) showing the β-barrel (teal), plug domain (orange). Residues R19 and D332 are likely involved in forming an ionic lock (R19-D332) that retains the Ton-box (magenta) sequestered within the β-barrel (A), as reported for BTUB_ECOLI (Zmyslowski et al., 2022), an *E. coli* vitamin B12 TBDR. (B) The CQNC motif is located at the tip of a prominent antiparallel β-hairpin, protruding toward the extracellular space from the plug domain. (C) The two cysteine thiol (sulfhydryl) side chains are close to each other (~3.1 Å; PAE = 1.54 Å), with the potential to form a disulfide bond in the oxidizing periplasmic environment during the protein's transit to the outer membrane. (E-G) AlphaFold3 models of the holoprotein with a bound $Mn^{2+}$ cation (ipTM = 0.84; pTM =0.90; mean pLDDT = 87.08) highlighting coordinating residues (C63, C66, E247, and H446) in tetrahedral geometry (G). (H) Boxplot analysis of donor-metal distances between $Mn^{2+}$ and the coordination donor atoms in Cys (median = 2.3245 ± 0.0259 median absolute deviation, $n$ = 8), Glu (2.08 ± 0.1572, $n$ = 53), and His (2.183 ± 0.2165, $n$ = 66) residues extracted from the MESPEUS database of coordination groups in proteins (Lin et al., 2024) with high-resolution structures ( ≤ 1.8 Å). The analysis reveals that the distances measured on the AF3 model (G) are very close to the median distances measured on crystal structures (H). Taken together, the high AF3 ipTM score, metal-donor distance measurements, and coordination geometry strongly suggest that residues C63, C66, E247, and H446 are directly involved in metal coordination. (I) AF3 model for the holoprotein with bound $Mn^{2+}$ predicted an allosteric conformational switch, increasing the distance between the R19 and D332 residues (4.5 Å; PAE = 2.5), releasing the predicted ionic lock, potentially allowing the N-terminal domain containing the Ton-box to extend into the periplasm (F), as demonstrated in BTUB_ECOLI (Zmyslowski et al., 2022). Abbreviations for AlphaFold metrics are explained at the end of the document.

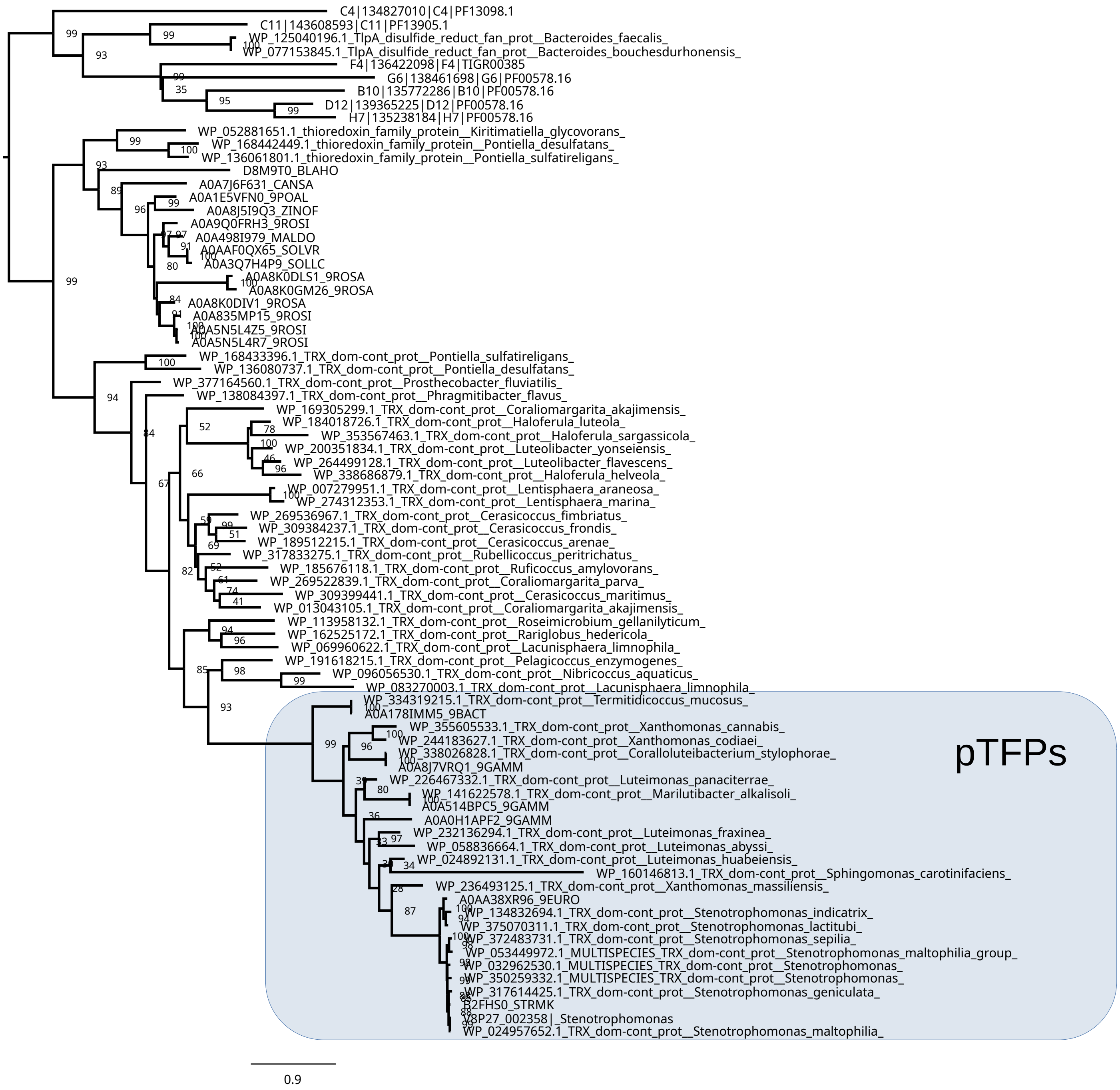


**Figure S2.** Maximum likelihood phylogeny estimated from 78 V8P27_002358 homologs collected from the NCBI RefSeq database (WP_ codes) via blastp, selected metagenomic sequences of thioredoxin superfamily proteins compiled by Nilewski et al. (7), and the UniProt Reference Proteomes v2025_01 identified by a hmmsearch through the HMMER portal using our pTFP.hmm profile HMM (see Methods). A highly supported clade (box) groups V8P27_002358 with other bacterial sequences belonging to the orders Lysobacterales (Gammaproteobacteria), Sphingomonadales (Alphaproteobacteria), and Opitutales (Verrucomicrobiota; Opitutia). The scale bar represents the number of expected substitutions per site under the best-fitting Q.pfam+I+R4 model chosen according to the Bayesian Information criterion, as implemented in IQ-Tree (Minh et al., 2020). Internal nodes are labeled with bootstrap support values computed by IQ-Tree with UFBoot.

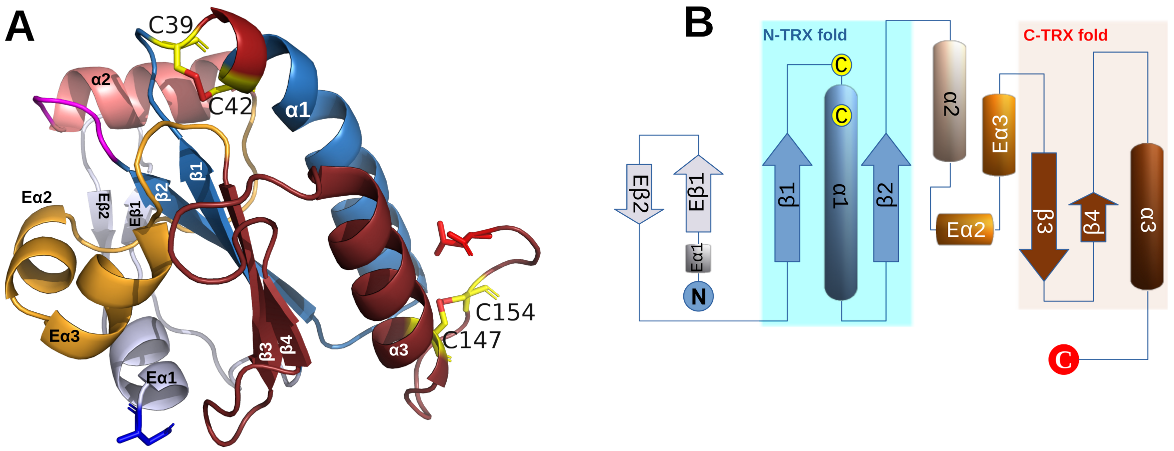


**Figure S3.** Structural models of the novel periplasmic thioredoxin-fold protein (pTFP) V8P27_002358 from *Stenotrophomonas maltophilia* Sm18 linked to the CQNC-type TBDRs shown in Fig. S1 by synteny and co-expression. (A) AlphaFold3 model (pTM = 0.93; mean pLDDT = 92.35) rendered with PyMOL (DeLano, 2004) and (B) 2D cartoon of V8P27_002358 showing canonical thioredoxin-fold (TRX) organization (light blue and ruby units). Fig. S2A presents the oxidized version (disulfide bridge in dark red) of the highly conserved CXXC motif of thioredoxins and thioredoxin-like proteins, which is found at the beginning of the α1 helix of the TRX fold (C39-C42). A second disulfide bridge is potentially formed between the C-terminal residues C147-C154. (B) The 2D topological cartoon of the mature V8P27_002358 protein highlights the TRX-fold elements and the position of the short extra N-terminal α-helix (Eα1) and β-strands (Eβ1 and Eβ2), and the two extra C-terminal α-helices (Eα2 and Eα3).

#### **AlphaFold metrics** of structural prediction confidence:

- **pLDDT (Predicted Local Distance Difference Test):** A per-residue measure (0–100) indicating how accurately AlphaFold predicted the local environment of a residue. Higher scores (typically > 70) indicate higher confidence in that region's structure.
- **pTM (Predicted Template Modeling score):** A metric (0–1) used to assess the overall accuracy of the predicted protein structure. Higher values (typically > 0,5) suggest the overall fold is similar to the true structure.
- **ipTM (Interface Predicted Template Modeling score):** A specific metric for protein complexes (multimers) that evaluates the quality of the interaction between different chains. It combines the accuracy of the interface with the overall structure.
- **PAE (Predicted Aligned Error):** A 2D matrix that estimates the positional error (in Ångströms) between two residues (*X* and *Y*) if the model was aligned on another part of the structure. It indicates confidence in the relative positions of domains or subunits.

#### Supplementary Referencences


Abramson, J., Adler, J., Dunger, J., Evans, R., Green, T., Pritzel, A., Ronneberger, O., Willmore, L., Ballard, A.J., Bambrick, J., Bodenstein, S.W., Evans, D.A., Hung, C.-C., O'Neill, M., Reiman, D., Tunyasuvunakool, K., Wu, Z., Žemgulytė, A., Arvaniti, E., Beattie, C., Bertolli, O., Bridgland, A., Cherepanov, A., Congreve, M., Cowen-Rivers, A.I., Cowie, A., Figurnov, M., Fuchs, F.B., Gladman, H., Jain, R., Khan, Y.A., Low, C.M.R., Perlin, K., Potapenko, A., Savy, P., Singh, S., Stecula, A., Thillaisundaram, A., Tong, C., Yakneen, S., Zhong, E.D., Zielinski, M., Žídek, A., Bapst, V., Kohli, P., Jaderberg, M., Hassabis, D., Jumper, J.M., 2024. Accurate structure prediction of biomolecular interactions with AlphaFold 3. Nature 1–3. https://doi.org/10.1038/s41586-024-07487-w

DeLano, W.L., 2004. Use of PyMOL as a communications tool for molecular science., in: Abstracts of Papers of the American Chemical Society. AMER CHEMICAL SOC 1155 16TH ST, NW, WASHINGTON, DC 20036 USA, pp. U313–U314.

Lin, G.-Y., Su, Y.-C., Huang, Y.L., Hsin, K.-Y., 2024. MESPEUS: a database of metal coordination groups in proteins. Nucleic Acids Res. 52, D483–D493. https://doi.org/10.1093/nar/gkad1009

Minh, B.Q., Schmidt, H.A., Chernomor, O., Schrempf, D., Woodhams, M.D., von Haeseler, A., Lanfear, R., 2020. IQ-TREE 2: New Models and Efficient Methods for Phylogenetic Inference in the Genomic Era. Mol. Biol. Evol. 37, 1530–1534. https://doi.org/10.1093/molbev/msaa015

Zmyslowski, A.M., Baxa, M.C., Gagnon, I.A., Sosnick, T.R., 2022. HDX-MS performed on BtuB in E. coli outer membranes delineates the luminal domain's allostery and unfolding upon B12 and TonB binding. Proc. Natl. Acad. Sci. U. S. A. 119, e2119436119. https://doi.org/10.1073/pnas.2119436119
